# The N-terminal and central domains of CoV-2 nsp1 play key functional roles in suppression of cellular gene expression and preservation of viral gene expression

**DOI:** 10.1101/2021.05.28.446204

**Authors:** Aaron S. Mendez, Michael Ly, Angélica M. González-Sánchez, Ella Hartenian, Nicholas T. Ingolia, Jamie H. Cate, Britt A. Glaunsinger

**Author notes:** equal contribution (Aaron S. Mendez, Michael Ly).

## Abstract

Nonstructural protein 1 (nsp1) is the first viral protein synthesized during coronavirus (CoV) infection and is a key virulence factor that dampens the innate immune response. It restricts cellular gene expression through a combination of inhibiting translation by blocking the mRNA entry channel of the 40S ribosomal subunit and by promoting mRNA degradation. We performed a detailed structure-guided mutational analysis of CoV-2 nsp1 coupled with *in vitro* and cell-based functional assays, revealing insight into how it coordinates these activities against host but not viral mRNA. We found that residues in the N-terminal and central regions of nsp1 not involved in docking into the 40S mRNA entry channel nonetheless stabilize its association with the ribosome and mRNA, thereby enhancing its restriction of host gene expression. These residues are also critical for the ability of mRNA containing the CoV-2 leader sequence to escape translational repression. Notably, we identify CoV-2 nsp1 mutants that gain the ability to repress translation of viral leader-containing transcripts. These data support a model in which viral mRNA binding functionally alters the association of nsp1 with the ribosome, which has implications for drug targeting and understanding how engineered or emerging mutations in CoV-2 nsp1 could attenuate the virus.

## Introduction

Viral infections frequently result in massive remodeling of the gene expression landscape within the cell, due to a combination of populating the cell with viral transcripts, inducing innate immune pathways and the activity of viral proteins that hijack or restrict key gene expression machinery. Protein synthesis is a focal point of control, as all viruses rely on cellular ribosomes for their protein synthesis and thus compete with endogenous mRNA for access to the translation machinery. A common viral strategy to shift translational resources towards viral mRNA is to restrict host gene expression, for example by inhibiting cap-dependent translation or encoding nucleases that degrade the cellular mRNA pool^1–4^. This phenotype, termed host shutoff, both increases viral transcript access to ribosomes and also promotes innate immune evasion^5^.

Host shutoff is a prominent feature of coronavirus (CoV) infection and has been shown to contribute significantly to suppression of innate immune responses in multiple pathogenic coronaviruses, including severe acute respiratory syndrome (SARS) CoV, Middle East respiratory syndrome (MERS) CoV and the pandemic CoV-2^6–8^. CoV-2-induced host shutoff is multi-faceted and involves inhibition of host mRNA splicing by the nonstructural protein (nsp) 16, restriction of cellular cytoplasmic mRNA accumulation and translation by nsp1 and disruption of protein secretion by nsp8 and nsp9^9–12^. Among these, the contribution and mechanism of action of nsp1 is best understood, in part because nsp1 was previously characterized as a key virulence factor in SARS CoV and related betacoronaviruses like mouse hepatitis virus (MHV)^13,14^. Indeed, an MHV mutant lacking functional nsp1 is severely attenuated in infected mice and mutation of nsp1 has thus been explored as a strategy for the development of a live attenuated vaccine for SARS CoV^14–16^.

Foundational work with SARS CoV nsp1 together with recent structural insights for the highly homologous CoV-2 nsp1 established that it engages in a bifunctional mechanism of shutoff of cytoplasmic mRNA that is unique amongst all characterized viral proteins^17–21^. Nsp1 binds directly to the 40S ribosomal subunit and positions its carboxyl-terminal (C-terminal) domain in the 40S mRNA entry channel, thereby blocking transcript access to the ribosome^22–25^. Structural studies revealed that the C-terminal helices of nsp1 dock within the entry channel through interactions with the RPS2, RPS3 and RPS30A ribosomal proteins as well as 18S rRNA^22–25^. These interactions are likely allosterically enhanced by the initiation factor eIF1, perhaps because eIF1 induces an ‘open head’ conformation of the 40S subunit mRNA entry channel that is favorable for nsp1 binding^26^. In addition to blocking translation, SARS CoV nsp1 also promotes cleavage of mRNA near the transcript 5’ end and CoV-2 nsp1 similarly reduces mRNA levels in cells^19–21,27^. Unlike other virally encoded host shutoff factors that deplete mRNA, nsp1 itself has no apparent ribonuclease activity and thus the mechanism underlying mRNA cleavage remains unknown^8,19–21^.

SARS CoV and CoV-2 nsp1 are 20 kilodalton (kDa) proteins with three general domains. The well-characterized helical C-terminal domain is connected to a globular amino terminal (N-terminal) domain through an unstructured central region. Mutations within the C-terminal domain that disrupt 18S ribosomal RNA (and thus 40S subunit) binding also block RNA cleavage, suggesting that mRNA cleavage is linked to translational repression^19–21^. Further, the identification of a double point mutant at the border of the N-terminal and central domains of SARS CoV nsp1 that disrupts mRNA cleavage but retains translational repression activity has led to the hypothesis that mRNA cleavage occurs subsequent to translational repression and is a functionally separable phenotype^21^. The structure of the N-terminal domain in isolation has been solved, but neither it nor the unstructured central region was resolved in the cryo-electron microscopy structures of CoV-2 nsp1 bound to the 40S ribosomal subunit and thus their role in host shutoff is unclear^28,29^.

Notably, CoV mRNAs contain a common 5’ leader sequence that protects them against nsp1-imposed host shutoff^20^. Studies with SARS-CoV nsp1 and CoV-2 nsp1 have implicated the N-terminal domain in binding the first stem loop (SL1) in the viral leader sequence to somehow facilitate the continued translation of viral mRNA^24,30,31^. Thus, deciphering the mechanistic contributions of domains outside of the C-terminal region in nsp1-induced host shutoff and viral escape will be key to ultimately understanding CoV virulence. Indeed, a recent CoV-2 genomic monitoring study reported a recurrent viral variant found in 37 countries containing an 11 amino acid (aa) deletion in the N-terminus of nsp1 that is associated with lower viral load, lower serum IFN-β and enrichment of less severe disease^32,33^.

Here, we define the functional contributions of each of the three nsp1 domains by performing a structure-function analysis of CoV-2 nsp1 using a combination of purified proteins and cell-based activity assays. Our results demonstrate that regions outside of the defined 40S interaction domain, including conserved residues in the N-terminal and central domains of nsp1 contribute to interactions with the 40S ribosomal subunit and with mRNA. Nsp1 mRNA binding and cleavage appear to occur only in the context of ribosome binding. While mRNA containing the CoV-2 leader sequence escapes repression by wild-type (WT) nsp1, we identify mutations in nsp1 that render these transcripts susceptible to potent translational inhibition. We hypothesize that an nsp1-40S-mRNA complex positions cellular mRNA for cleavage when the mRNA cannot engage the ribosome entry tunnel, while viral mRNA escapes cleavage because its association with nsp1 triggers remodeling of the complex to enable translation. However, if viral mRNA is not properly engaged by nsp1, it becomes susceptible to repression. It is therefore likely that mutations within the N-terminal domain of nsp1 could decrease CoV-2 virulence both by decreasing host shutoff and by restricting viral gene expression. In this regard, drugs that target the interaction between the nsp1 N-terminus and mRNA may be good candidates for antiviral therapy.

## Results

### CoV-2 nsp1 promotes translational shutoff and mRNA decay

SARS-CoV nsp1 and CoV-2 nsp1 are 84.4% identical at the amino acid level, including conservation of residues characterized in SARS-CoV nsp1 as required for nsp1 binding to the 40S subunit (K164/H165) and for promoting mRNA turnover (R124/K125) (Figure S1A)^19,21,34^. To confirm that CoV-2 nsp1 functions in a similarly bifunctional manner as SARS-CoV nsp1, we measured its translation repression and RNA cleavage activity *in vitro* and in cells. We assume that the changes in protein output in these assays are a combined reflection of changes to translational efficiency and mRNA abundance. As expected, when added to HEK293T *in vitro* translation extracts, purified CoV-2 nsp1 markedly suppressed translation of a nanoluciferase reporter containing the 5’ UTR of human β-globin (HBB-nLuc) in a dose dependent manner (Figure 1A). Introducing K164A/H165A mutations, which inhibit 40S binding, also abolishes translation inhibition (Figure 1A), while mutant R124A/K125A largely retained the ability to suppress nLuc translation (Figure 1A). Similar results were obtained with rabbit reticulocyte lysates (Figure S1B). Nsp1 also induced HBB-nLuc mRNA cleavage in translation extracts, as measured by primer extension assays (Figure 1B). Cleavage required nsp1 binding to ribosomes, as the mRNA was uncleaved in the absence of translation extracts or by the mutant K164A/H165A (Figure 1B). Nsp1 R124A/K125A showed reduced mRNA cleavage relative to WT protein, confirming that these residues within the central nsp1 domain are involved in its mRNA turnover activity (Figure 1B).

**Figure 1.**
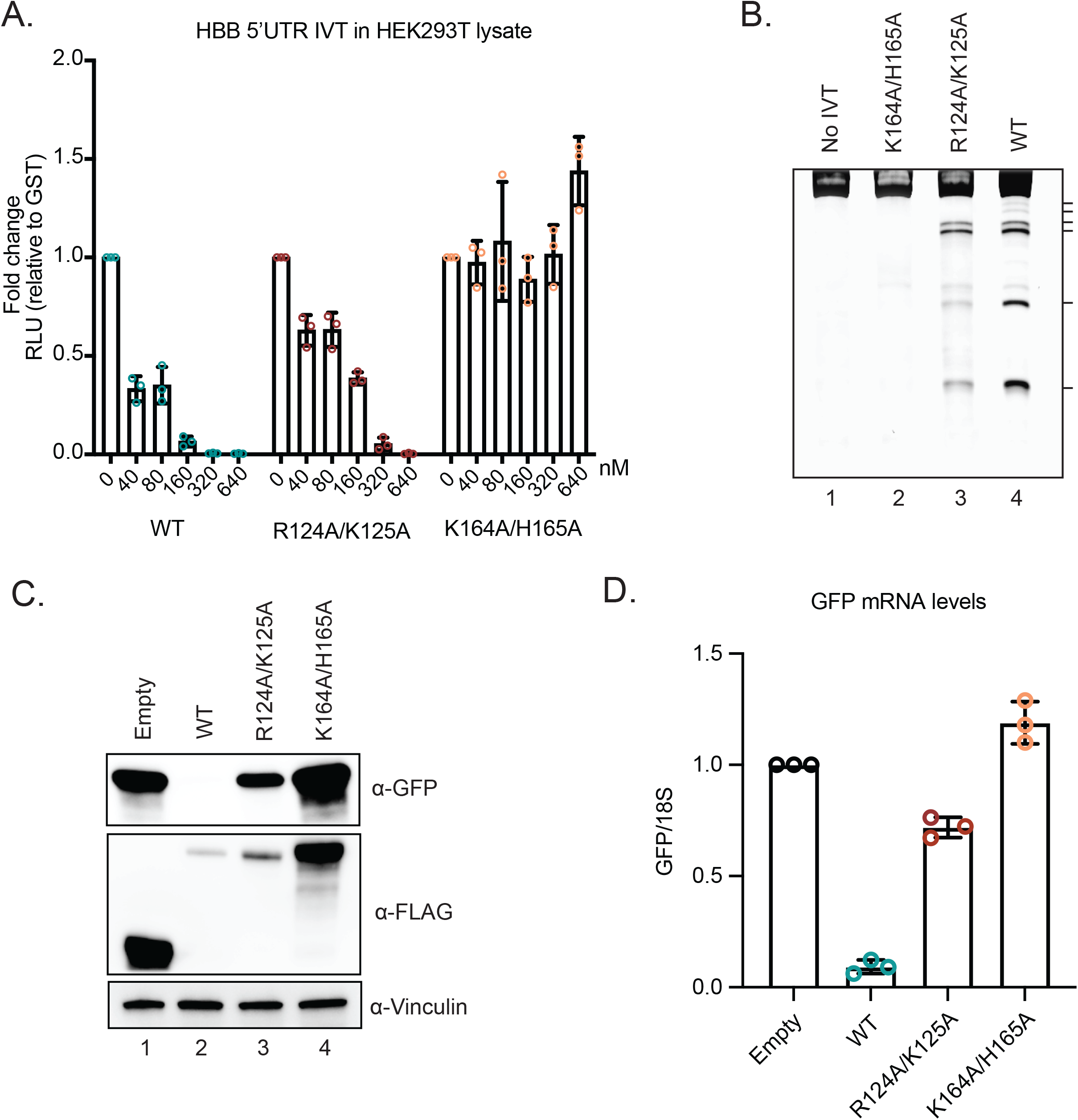
CoV-2 nsp1 promotes translational suppression and mRNA decay in vitro and in cells. (A) HBB-nLuc reporter RNA was incubated with HEK293T translation extracts alone or in the presence of increasing concentrations of purified WT, R124A/K125A or K164A/H165A nsp1. Translation of the reporter was then evaluated by luciferase assay and normalized to a GST protein control. (B) A primer extension assay was used to measure cleavage of the HBB-nLuc RNA in the presence of purified WT or mutant nsp1. Lane 1 (no IVT) shows nsp1 and HBB-nLuc incubation in primer extension buffer only whereas lanes 2-4 show reactions incubated in the presence of translation extracts. Hash marks denote cleavage intermediates. (C-D) HEK293T cells were transfected with a GFP reporter plasmid alone or together with the indicated nsp1 expressing plasmids and then harvested for protein or RNA. GFP and nsp1 protein levels were measured by α-GFP and α-FLAG western blots, respectively, with vinculin used as a protein loading control (C). GFP mRNA was quantified by RT-qPCR and normalized to 18S rRNA, with the level of GFP mRNA in cells lacking nsp1 then set to 1 (D).

Finally, we monitored CoV-2 nsp1 activity in cells by co-transfecting HEK293T cells with plasmids expressing WT or mutant nsp1 together with a GFP reporter plasmid, then evaluating levels of GFP protein by western blot and GFP mRNA by RT-qPCR. WT nsp1 prevented GFP protein expression and caused a 40-fold reduction of GFP transcript levels (Figure 1C-D). In line with the *in vitro* assays, the K164A/H165A mutant had no impact on GFP protein or mRNA, while the R124A/K125A mutant caused a modest reduction in GFP protein and a 2-fold reduction of GFP mRNA (Figure 1C-D). Note that nsp1 can target its own transcript for repression, so in these and subsequent cell-based assays the level of nsp1 protein generally reflects its activity (i.e. WT nsp1 accumulates to lower levels than functionally defective mutants).

### The N-terminal and central domains of nsp1 are required for host shutoff

The above results and recent published data have confirmed the essential role of the C-terminal domain of CoV-2 nsp1 in translational suppression and have shown that residues R124 and K125 in the central domain are important for promoting efficient mRNA decay. However, little is known about the contribution of the N-terminal domain to host shutoff activity or the mechanistic role of the central domain. To address these questions, we first constructed a series of deletion mutants in which we removed either the N-terminal domain (Δ 1-117) or the C-terminal and central domains (Δ 118-180) (Figure 2A). We also tested whether the region containing the R124/K125 ‘RNA destabilization’ residues had a sequence-dependent role or a sequence-independent spacing role by deleting residues encompassing this region (Δ122-130) and then replacing them with a size-matched glycine linker (G-linker). We evaluated these mutants using the cell-based assays described above to measure protein and mRNA levels of the GFP reporter. The WT and mutant nsp1 proteins were tagged with a 3xFLAG-Halo tag for detection, which we confirmed does not interfere with nsp1 function (Figure S2A).

**Figure 2.**
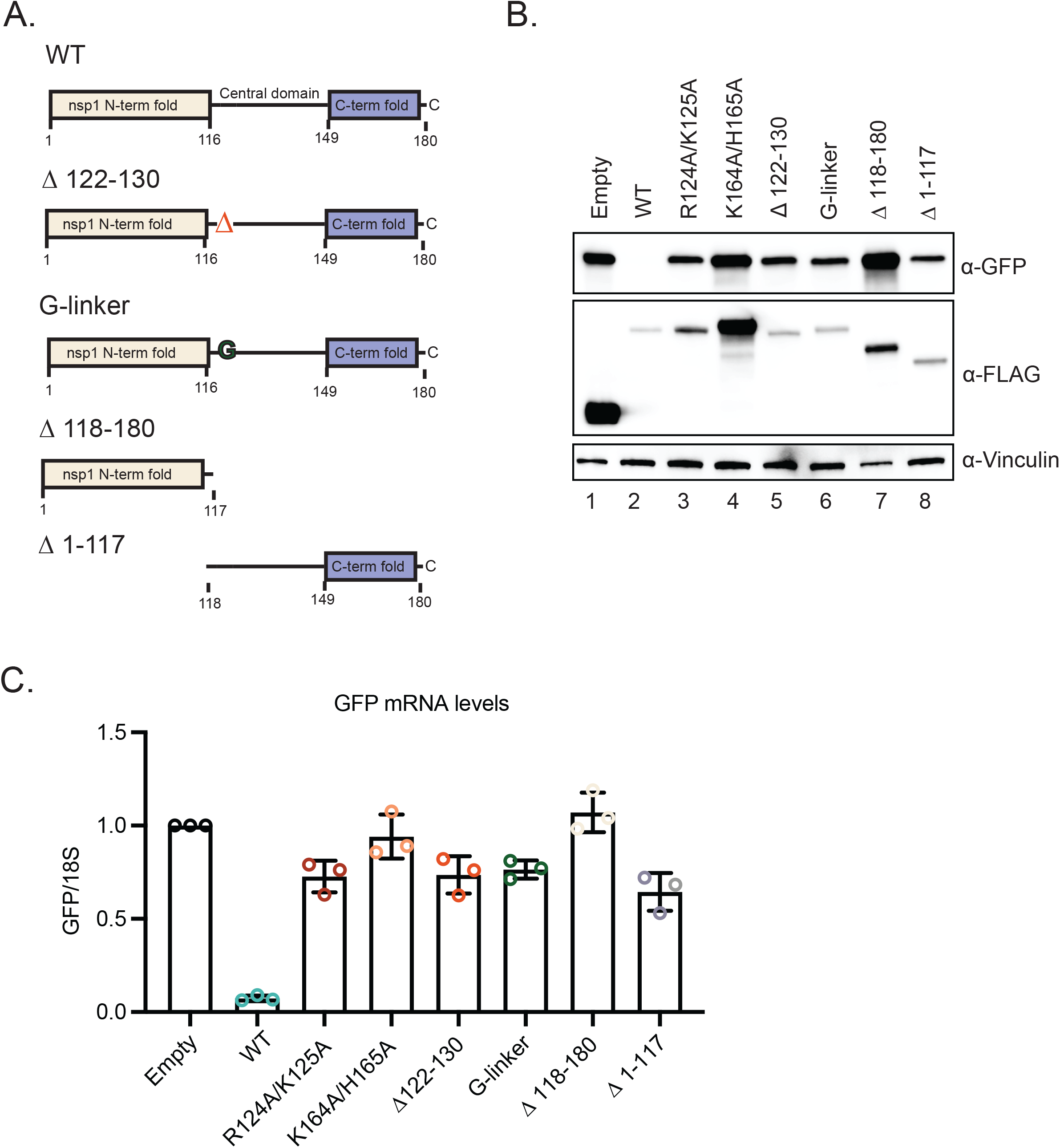
The N-terminal and central domains of nsp1 are required for translational suppression and mRNA depletion. (A) Schematic of the 3xFLAG-Halo-tagged versions of WT and mutant nsp1. Amino acids 122-130 encompass the RNA destabilization domain, which was either deleted (Δ122-130) or replaced with a size matched glycine linker (G-linker). Mutant Δ118-180 lacks the central and C-terminal domains, while Δ1-117 lacks the N-terminal domain. (B-C) HEK293T cells were transfected with a GFP reporter plasmid alone or together with plasmids containing WT or the indicated mutant nsp1 and then harvested for protein or RNA. GFP and nsp1 protein levels were measured by α-GFP and α-FLAG western blots, respectively, with vinculin used as a protein loading control (B). GFP mRNA was quantified by RT-qPCR and normalized to 18S rRNA, with the level of GFP mRNA in cells lacking nsp1 then set to 1 (C).

Notably, we found that in the absence of its N-terminal domain, nsp1-induced shutoff of both GFP protein and mRNA to reduced levels similar to the R124A/K125A mutant (Figure 2B-C). The N-terminal domain alone showed no translational suppression or mRNA turnover activity, which is not surprising given that it lacks the K164/H165 residues essential for 40S subunit binding (Figure 2B-C). The Δ122-130 and G-linker mutants behaved similarly to the R124A/K125A point mutant, suggesting that this region has a more specific function and is not simply a flexible spacer separating the N-terminal and C-terminal domains (Figure 2B-C). Together, these data establish that residues within the N-terminus of nsp1 play key roles in host shutoff.

### Residue R99 in the N-terminal domain of nsp1 contributes to its mRNA destabilization function and translational shutoff

We next sought to better define which residues within the N-terminal domain are required for suppressing gene expression. Guided by existing structural data for the N-terminus^27,28^, we mutated a series of conserved, surface exposed and charged residues (E36A/E37A, E55A/E57A/K58A, R99A and R119A/K120A) (Figure S3A). The mutants were then screened using our GFP reporter assay alongside the R124A/K125A and K164A/H165A controls to determine the effects on protein translation and mRNA decay. Most of the N-terminal domain mutants behaved similarly to WT nsp1 in their ability to suppress GFP protein and mRNA levels, although E36A/E37A was modestly impaired for mRNA depletion (Figure 3A-B). The one exception was mutant R99A, which displayed a moderate defect in restricting GFP protein accumulation and a severe defect in mRNA targeting (Figure 3A-B).

**Figure 3.**
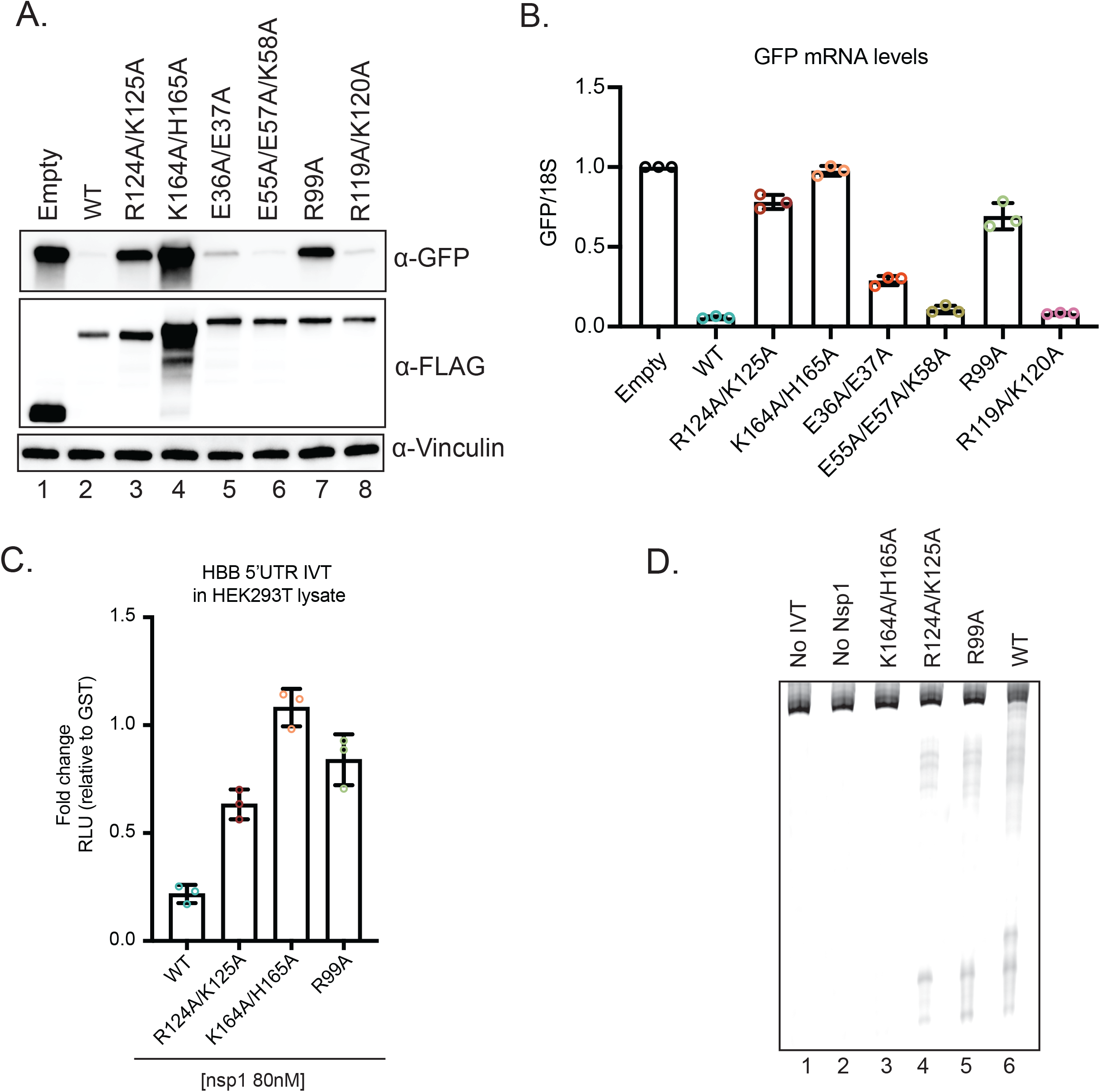
Residue R99 located in the N-terminal domain plays key roles in nsp1-induced host shutoff. (A-B) HEK293T cells were transfected with a GFP reporter plasmid alone or together with plasmids containing WT or the indicated mutant nsp1 and then harvested for protein or RNA. GFP and nsp1 protein levels were measured by α-GFP and α-FLAG western blots, respectively, with vinculin used as a protein loading control (*A*). GFP mRNA was quantified by RT-qPCR and normalized to 18S rRNA, with the level of GFP mRNA in cells lacking nsp1 then set to 1 (*B*). (C) HBB-nLuc reporter RNA was incubated with HEK293T translation extracts in the presence of 80 nM of purified WT or the indicated mutant nsp1 protein. Translation of the reporter was then evaluated by luciferase assay and normalized to levels from lysates incubated with 80 nM of a control GST protein. (D) Primer extension assay to measure degradation of the HBB-nLuc RNA in the presence and absence of purified WT or mutant nsp1. Lanes 1 and 2 are controls lacking translation extract (no IVT) or nsp1, respectively.

The R99A defect was also observed when we purified the protein and tested its activity in the HEK293T translation extracts. Compared to WT nsp1, which suppressed translation of HBB-nLuc 17-fold, the R99A mutant suppressed translation by only 1.8-fold, similar to the 2-fold suppression seen with the R124A/K125A mutant (Figure 3C). This defect persisted even at high concentrations of the R99A protein (Figure S3C). Primer extensions revealed that R99A showed a modest reduction in nsp1-induced RNA cleavage, similar to R124A/K125A (Figure 3D). We note that while R99A, R124A/K125A and K164A/H165A mutants all show pronounced defects in mRNA depletion in cells, only the 40S binding mutant strongly disrupts RNA cleavage in the *in vitro* assay.

### The nsp1 N-terminus and central domain are involved in ribosome binding

To identify the mechanism(s) underlying the defects of the various CoV-2 nsp1 domain mutants, we first evaluated their ability to bind the 40S ribosomal subunit both in cells and using quantitative *in vitro* measurements. Binding in cells was evaluated by immunoprecipitation (IP) of nsp1 followed by western blotting to monitor co-purification of the 40S subunit proteins RPS2, RPS3, RPS24 and RACK1, which are enriched as nsp1 interaction partners in published mass spectrometry data sets^10,35^ (Figure 4A). Notably, all mutants we tested in this assay showed reduced levels of ribosomal protein interactions, including the point mutations R124A/K125A, E36A/E37A and (to a lesser extent) R99A located in the N-terminus, and the deletion mutant Δ1-117 lacking the N-terminal domain (Figure 4A). As expected, no interaction with 40S proteins was observed in the absence of the C-terminal domain (Δ118-180) or with the K164A/H165A 40S binding mutant.

**Figure 4.**
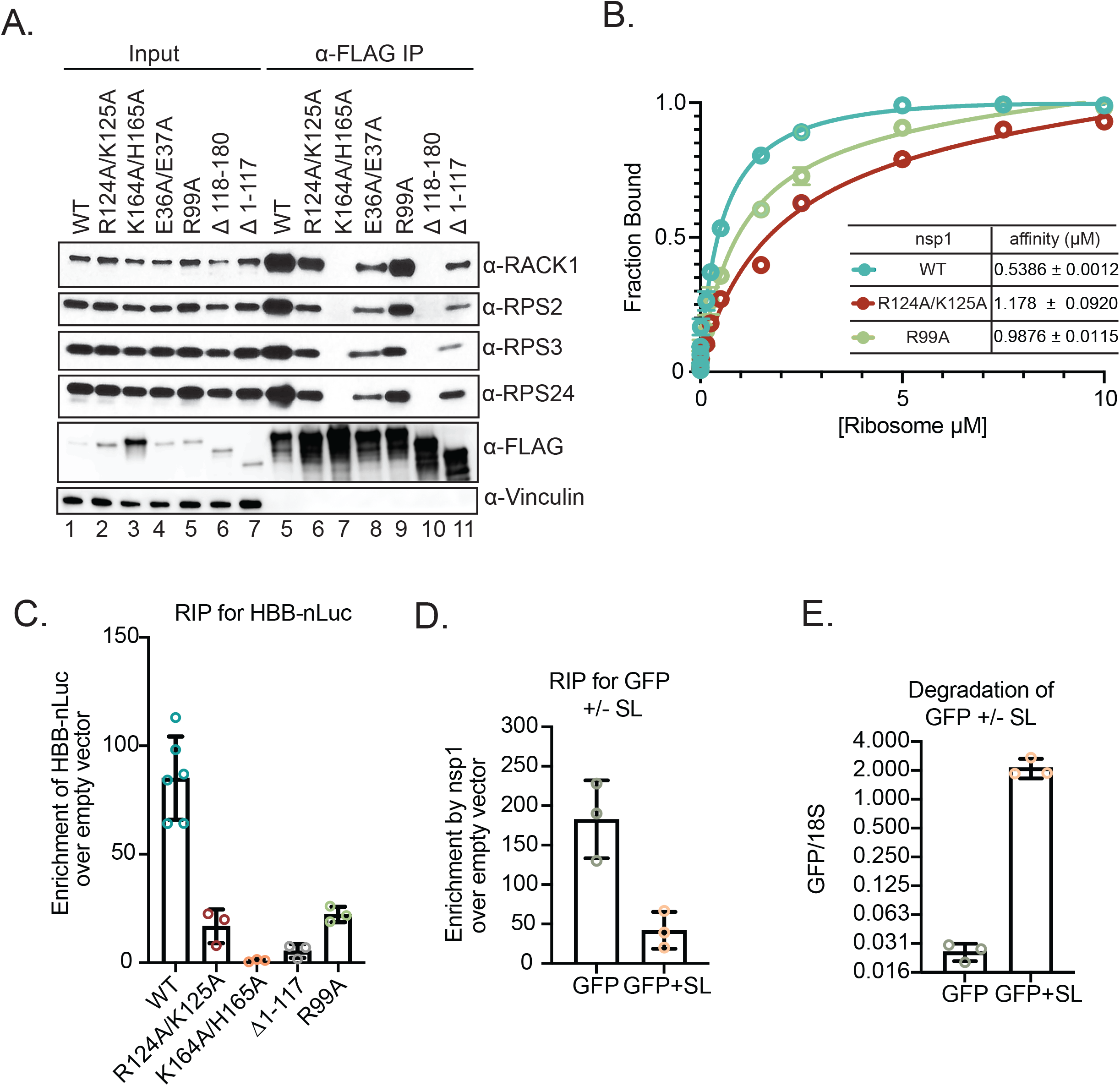
Nsp1 N-terminal and central domain mutants are defective for ribosome and mRNA binding. (A) HEK293T cells were transfected with plasmids expressing WT or the indicated mutant 3x-FLAG-Halo-tagged nsp1. Nsp1 was immunoprecipitated (IP) using α-FLAG beads and co-IP of ribosomal proteins RACK1, RPS2, RPS3 and RPS24 was monitored by western blotting, with vinculin serving as a loading control. Input lanes contain 1/10 of the amount of protein used for the IPs. (B) Equilibrium binding measurements of fluorescently labeled WT, (blue), R124A,K125A (red) and R99A (green) nsp1 to purified ribosomes. Data represent a total of 3 biological replicates. (C) HEK293T cells were co-transfected with HBB-nLuc and either a control plasmid or the indicated 3X-FLAG-Halo-tagged nsp1 constructs. Nsp1 was immunoprecipitated using α-FLAG beads, whereupon the co-immunoprecipitating RNAs were extracted and nLuc mRNA was quantified by RT-qPCR. The mRNA values were then normalized to the values obtained from the empty vector control. (D) HEK293T cells were co-transfected with a 3X-FLAG-Halo-tagged nsp1 plasmid or empty vector control, together with a plasmid expressing either GFP with a 5’ stem loop (GFP+SL) or a control GFP lacking the stem loop (GFP). Nsp1 was immunoprecipitated using α-FLAG beads, whereupon the co-immunoprecipitating GFP+SL or GFP mRNAs were quantified by RT-qPCR. The mRNA values were then normalized to those obtained from the empty vector control. (E) The levels of GFP+SL and GFP mRNA present in the input samples from (C) were quantified by RT-qPCR and normalized to 18S rRNA, with the level of GFP mRNA in cells lacking nsp1 (empty vector control) set to 1.

We next quantified ribosome affinity directly using fluorescent polarization (FP) equilibrium binding experiments in the presence of increasing concentration of purified ribosomes. We site-specifically labeled WT nsp1 and nsp1 mutants at an engineered internal position with a TAMRA dye for these ribosome binding measurements. Raw polarization units confirmed minimal binding of the K164A/H165A to ribosomes compared to WT nsp1, which bound with a dissociation constant (Kd) of 0.5386 μM (Figure S4A). In agreement with the reduced co-purification of 40S proteins from cells, R124A/K125A and R99A showed a 2-fold and a 1.6-fold increase in their Kd values, respectively (Figure 4B). Thus, while the C-terminal domain is critical for ribosome binding, both the N-terminal and central regions contribute to the stability of the interaction.

### Multiple nsp1 domains contribute to binding mRNA in association with the ribosome

In addition to their contribution to ribosome binding, we also considered the possibility that the nsp1 N-terminal and central domains may be involved in binding mRNA. We were unable to detect a direct interaction between purified nsp1 and RNA *in vitro* (data not shown), perhaps suggesting that nsp1 only binds RNA in the context of the 40S ribosomal subunit in cells. To test this, we evaluated the ability of WT and mutant nsp1 to bind the HBB-nLuc reporter mRNA in transfected HEK293T cells using RNA immunoprecipitation (RIP) experiments. Indeed, we observed an 85-fold enrichment of HBB-nLuc mRNA upon IP of the FLAG-tagged WT nsp1, but no binding to the reporter by the 40S binding mutant K164A/H165A and a 15-fold decrease in binding by the nsp1 Δ1-117 mutant lacking the N-terminal domain (Figure 4C and Figure S5A). RIP experiments with the R124A/K125A and R99A point mutants showed a 7-fold and 2.5-fold decrease in mRNA binding, respectively (Figure 4C). Importantly, RT-qPCR measurements of copurifying 18S rRNA in each RIP correlated with the level of 40S binding seen by western blotting (Figure S5B), suggesting that the observed nsp1-mRNA interactions are likely to be bridged by the 40S ribosomal subunit.

To more directly test whether an mRNA must be bound to 40S in order to associate with nsp1, we performed RIPs to evaluate the ability of FLAG-nsp1 to associate with a GFP mRNA containing a cap-proximal stem loop structure (GFP+SL) that blocks 40S binding and thus cannot be translated (Figure S6)^36,37^. Indeed, nsp1 binding to the GFP+SL was reduced >5-fold relative to the translation-competent version of GFP and nsp1 was incapable of degrading GFP+SL (Figure 4D-E). Collectively, these data suggest that the nsp1 N-terminus contributes to a stable nsp1-mRNA-40S subunit complex, which is needed to efficiently induce mRNA cleavage.

### CoV-2 leader sequence-mediated escape requires nsp1 residues R124/R125 and R99

The 5’ leader sequence of SARS-CoV and CoV-2 interact with nsp1 to somehow protect viral transcripts from its cleavage activity^20,24,30^. To evaluate the contribution of the nsp1 domains towards CoV-2 leader sequence binding, we generated an nLuc reporter mRNA containing the CoV-2 leader sequence at its 5’ end (CoV2L-nLuc). As expected, RIPs with WT FLAG-nsp1 significantly enriched for CoV2L-nLuc over empty vector control (Figure S7), whereas the 40S binding mutant K164A/H165A showed no enrichment for the CoV2L-nLuc reporter (Figure 5A). Similar to the results with the mRNA lacking the CoV-2 leader, the truncation mutant lacking the N-terminal domain (Δ1-117), the R124A/K125A and the R99A mutants all displayed reduced binding to the CoV-2 leader mRNA (a 32-fold, 20-fold and 10-fold reduction, respectively) (Figure 5A). Thus, like cellular mRNA, the transcript containing the CoV-2 leader does not readily associate with nsp1 in the absence of 40S binding.

**Figure 5:**
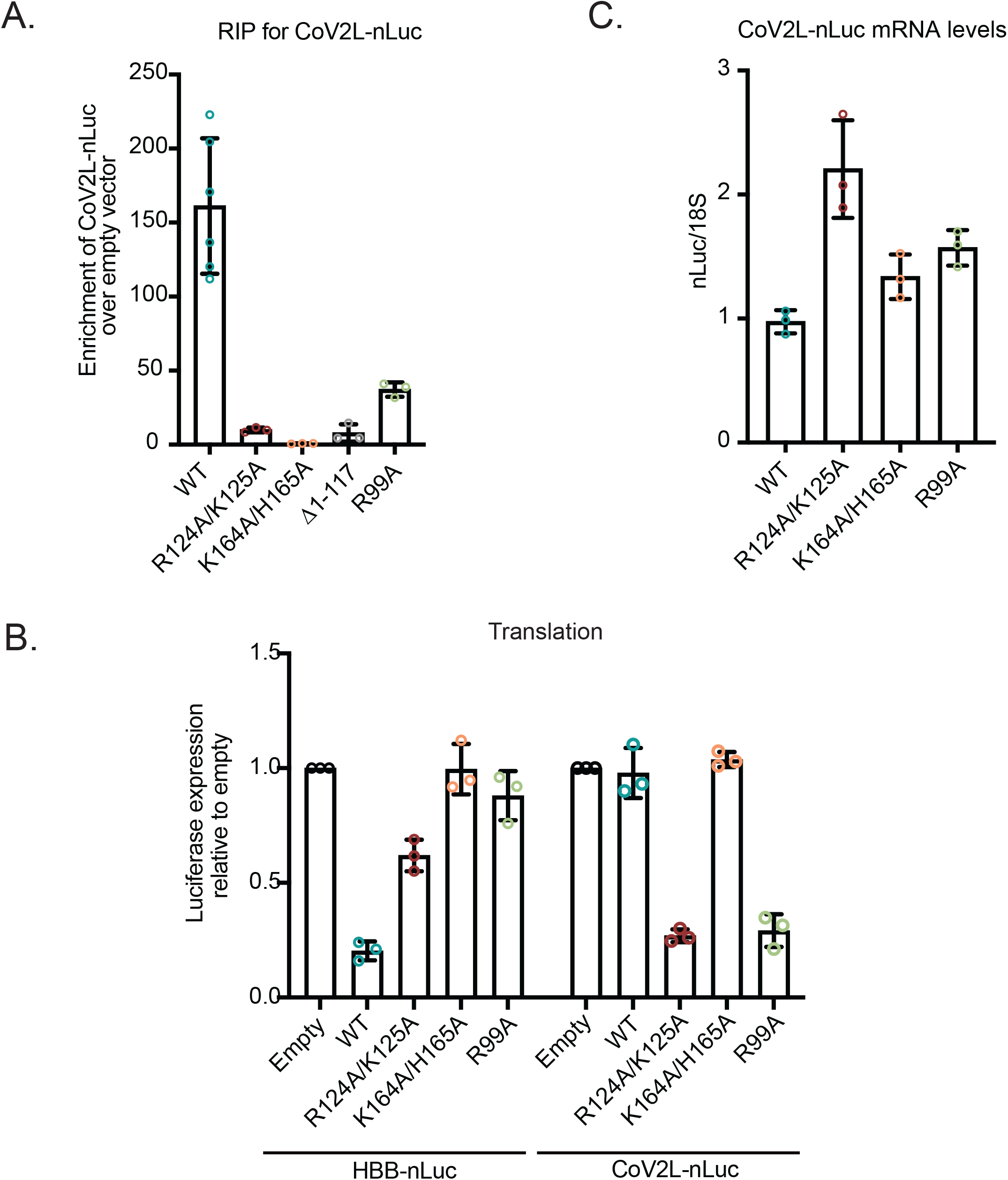
Protection from translational repression conferred by the CoV-2-leader sequence is selectively eliminated by nsp1 N-terminal and central domain mutants. (A) HEK293T cells were co-transfected with a plasmid expressing CoV-2 leader-nLuc and either a control plasmid or the indicated 3X-FLAG-Halo-tagged nsp1 construct. Nsp1 was immunoprecipitated using α-FLAG beads, whereupon the co-immunoprecipitating RNAs were quantified by RT-qPCR. The mRNA values were then normalized to the mRNA values obtained from the empty vector control. (B) HEK293T cells were transfected with either HBB-nLuc or CoV2L-nLuc together with control empty vector or the indicated nsp1 construct. Translation of HBB-nLuc or CoV2L-nLuc was measured by luciferase assay and the fold-change in luciferase activity was calculated relative to the empty vector control. (C) CoV2L-nLuc mRNA was quantified from the above experiment by RT-qPCR and normalized to 18S rRNA, with the level of CoV2L-nLuc mRNA in cells lacking nsp1 then set to 1.

Although the N-terminal and central domains of nsp1 participate in binding both cellular and viral mRNA, the consequences of this binding are presumably different. Indeed, the CoV2L-nLuc mRNA remained fully translationally competent in the presence of nsp1, under conditions that suppressed translation of HBB-nLuc (Figure 5B-C). Remarkably, however, both the R124A/K125A and R99A mutants gained the ability to translationally repress CoV2L-nLuc, although they remained impaired for translational suppression of HBB-nLuc (Figure 5B). Nsp1 K164A/H165A had no impact on translation of either reporter, as expected (Figure 5B).

Finally, we performed RT-qPCR to determine whether translational suppression of CoV2L-nLuc also resulted in degradation of the mRNA. Surprisingly, there was no decrease in CoV2L-nLuc mRNA abundance in the presence of these mutants, indicating that the decrease in protein expression was solely due to translational repression (Figure 5C). Collectively, these data show that the interaction of the viral leader sequence with nsp1 residues R124, K125 and R99 in an mRNA-nsp1-40S subunit ternary complex is critical for mediating escape from translational suppression, underscoring their functional importance for both host shutoff and efficient viral gene expression.

## Discussion

Virus-induced host shutoff has profound impacts on viral pathogenesis, as has been demonstrated for pathogens ranging from influenza virus to coronavirus to herpesviruses^14,38–41^. The mechanisms by which viral transcripts retain robust expression during host shutoff are generally tailored to the specific host shutoff strategy and include the use of alternative RNA processing or ribosome recruitment mechanisms, as well as kinetic regulation of the host shutoff factor^42–44^. Despite numerous studies on the dual translational repression and mRNA cleavage functions of nsp1, significant knowledge gaps exist for how it mechanistically coordinates both activities against cellular mRNA while sparing viral transcripts. Here, we report the most detailed structure-guided mutational analysis to date of nsp1. Using a combination of *in vitro* and cell-based assays to measure translation, ribosome binding, RNA binding and mRNA degradation by WT and mutant CoV-2 nsp1, we find that its host shutoff activities all appear to occur in the context of an nsp1-mRNA-40S complex. We also made two additional notable new findings that are summarized in the model shown in Figure 6. First, the CoV-2 nsp1 N-terminus and neighboring residues that do not contact the mRNA entry channel nonetheless stabilize the nsp1-40S subunit interaction and in doing so enhance its host shutoff functions. Second, we identify specific residues in nsp1 whose mutation abrogates the translational escape of viral leader-containing mRNA. Collectively, these findings provide insight into the functional contribution of nsp1 regions outside of the 40S ‘docking’ domain and have implications for understanding why alterations to these regions may impair viral pathogenesis.

**Figure 6.**
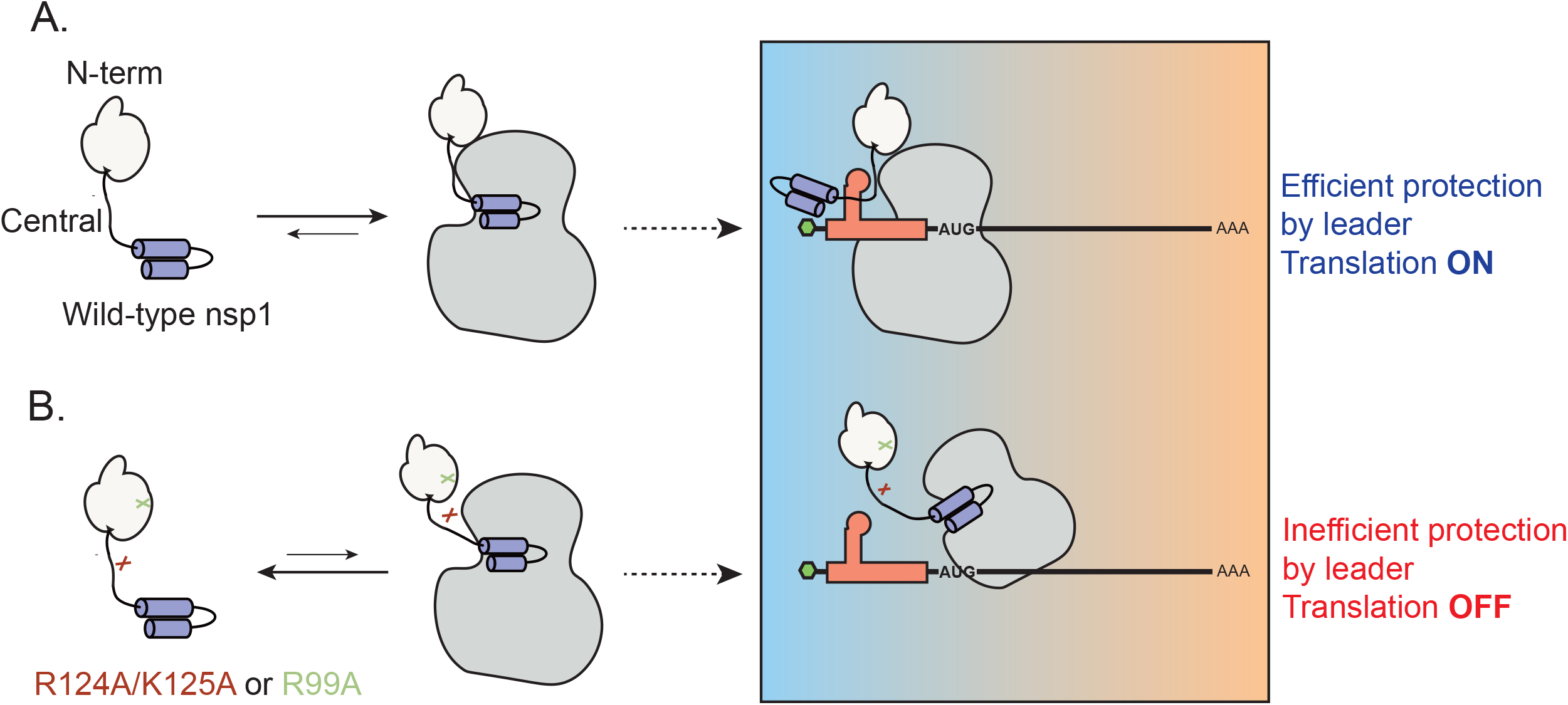
Model for how the N-terminal and central domains of nsp1 are critical for 40S ribosome association and preservation of leader-containing transcripts. (A) All three nsp1 domains contribute to its interaction with the 40S ribosome. While the C-terminal domain interjects into the mRNA entry channel of the ribosome to block mRNA access, the N-terminal and central domains stabilize the interaction. When cellular mRNA encounters an nsp1-bound ribosome, it is translationally blocked and undergoes degradation. However, mRNA containing the CoV-2 leader sequence engages the N-terminal and central domains of 40S-bound nsp1 in a manner involving nsp1 residues R124, K125, and R99, leading to relief from translational repression. (B) Nsp1 mutants R124A/K125A and R99A have reduced affinity for the 40S ribosome, which alleviates translational repression of cellular transcripts. However, CoV-2 leader-containing transcripts instead become translationally repressed, perhaps due to a ‘nonproductive’ interaction with nsp1 in the absence of proper engagement with residues R124/K125 or R99.

Mutation of CoV-2 nsp1 N-terminal and central domain residues R99, R124 and K125 compromise its binding to the 40S ribosomal subunit, as observed by reduced association with ribosomal proteins and 18S rRNA in IP experiments, as well as increased dissociation constants with purified ribosomes. While the nsp1 C-terminal domain docks in the 40S mRNA entry channel, if and where these other nsp1 domain interactions occur on the ribosome is unknown. It is possible that mRNA stabilizes the nsp1-40S complex through interactions with the N-terminal and central domains, as nsp1 mRNA binding decreases in accordance with reduced nsp1-40S binding. In this regard, a stable interaction between nsp1 and mRNA presumably does not occur prior to 40S subunit binding, as shown by the markedly reduced nsp1 binding to an mRNA with a 40S blocking sequence. Also, no cellular or CoV-2 leader sequence mRNA binding was detected using nsp1 K164A/H165, which fails to engage the 40S subunit. However, recent *in vitro* data also argue against nsp1 binding ribosomes that have already engaged mRNA in their entry channel, or eIF3j-bound ribosomes that have a ‘closed head’ conformation^25,26^. Thus, mRNA binding by nsp1 presumably occurs subsequent to nsp1-40S binding and/or with mRNA that is associated with the 40S subunit but has not yet engaged the entry channel. Cellular mRNA binding could either occur indirectly via bridging interactions with the 40S subunit or nsp1 could possess weak intrinsic mRNA binding activity that is significantly stabilized by 40S binding.

Our results indicate that weakening the nsp1-mRNA-40S subunit interaction rescues cellular mRNA from degradation, as does adding a 40S subunit blocking sequence to the mRNA. How the nsp1-mRNA interaction is stabilized by the 40S subunit and how this interaction leads to cleavage of the mRNA are open central questions. Given that the nsp1 C-terminal domain occludes the 40S subunit mRNA entry channel, the mRNA present in the tripartite nsp1-mRNA-40S subunit complex is presumably held elsewhere on the ribosome, perhaps through interactions with translation initiation factors. This notion is supported by the observation that mRNAs containing certain internal ribosome entry site (IRES) elements can escape cleavage by SARS-CoV nsp1, depending on their initiation factor requirements^18,19,21,45^. For example, mRNAs bearing a cricket paralysis virus IRES (which recruits the 40S ribosomal subunit in the absence of any initiation factors) or a hepatitis C virus IRES are not cleaved, whereas mRNAs with picornavirus type I and type II IRES elements are susceptible to SARS-CoV nsp1-induced cleavage^19,21,31^. In the future, it could be informative to test whether susceptible versus cleaved IRES elements have different abilities to bind the nsp1-40S complex.

Unlike cellular mRNA, viral mRNA remains robustly translated in the presence of nsp1^6,30,33,46^. As shown for both SARS-CoV and CoV-2, preservation of viral gene expression in the presence of nsp1 requires a conserved stem loop (termed SL1) within the 5’ leader sequence present on viral mRNAs^6,24,30,31^. Two main models have been put forth to explain viral mRNA escape. One posits that nsp1 can target both cellular and viral transcripts equally, but leader-containing transcripts have higher intrinsic translational efficiency than cellular mRNA and can thus preferentially engage free 40S ribosomes^22,23,25,26^. The other model proposes that viral mRNA is directly refractory to suppression by nsp1, perhaps because it interacts with nsp1 in a manner that causes an allosteric change in nsp1 that removes its C-terminal domain from the 40S entry channel^24,30,31,47^.

Although we agree that viral mRNA may have intrinsically high translational efficiency, several of our observations favor the second model. First, we found that mRNA bearing the CoV-2 leader was readily associated with nsp1-40S subunit complexes but was not subject to degradation. Even if a fraction of the CoV-2 leader mRNA was more efficiently engaged by non-nsp1 bound 40S subunits, in the absence of protection we would still have expected to see cleavage of those bound by the nsp1-40S ribosomes. Second, and more telling, are the observations that specific mutations within and bordering the N-terminal domain of nsp1 (R99A, R124A/K125A) have negative rather than neutral effects on translation of CoV-2 leader mRNA. Both of these mutants have weaker association with ribosomes and mRNA, and thus would be expected to free up even more 40S ribosomes for efficient translation of viral transcripts. Instead, R99A and R124A/K125A gain the ability to potently suppress translation of CoV-2 leader-containing transcripts. We therefore propose that these residues are crucial for viral leader sequence-triggered conformational changes to nsp1 that enable viral mRNA translation. Their mutation may either prevent proper binding to viral mRNA and/or ‘lock’ nsp1 in a translationally repressive conformation. Intriguingly, these mutants translationally repress CoV-2 leader mRNA translation without reducing its mRNA abundance, which may have implications for understanding how mRNA cleavage is activated. Notably, the above model is supported by the observation that mutating the central domain residue R124 in SARS-CoV nsp1 reduces viral gene expression, suggesting conservation across betacoronaviruses in the functional role for regions outside of the C-terminus in promoting viral escape^29^.

The finding that nsp1 point mutants like R99A can abrogate translational protection of viral leader-containing mRNA while having reduced host shutoff activity against cellular mRNA has important implications for understanding the role of nsp1 in pathogenesis. *In vivo* experiments with the model betacoronavirus MHV showed that deletion of residues within the nsp1 central domain results in reduced viral load and heightened survival rates and this virus showed promise as a live attenuated vaccine platform^16^. Additionally, mutation of the nsp1 R124 residue in the SARS-CoV replicon system decreased viral gene expression and replication^30^. It is therefore possible that specific nsp1 mutations such as R99, R124 and R125 could impair viral pathogenesis as a consequence of translational suppression of viral transcripts in addition to impaired shutoff of host genes such as those involved in interferon signaling. It will be of interest to determine whether this underlies the lower viral load and decreased pathogenesis reported for the recurrent CoV-2 variant that contains an 11 aa nsp1 N-terminal domain deletion, or other variants that may emerge with changes to these regions of nsp1^33^. Likewise, small molecule drugs that phenocopy these mutations could serve as therapies or prophylactics for viral infection.

## Materials and Methods

### Cloning and mutagenesis

All plasmids generated for this study have been deposited in Addgene (*pending*). Sequences for the oligos and gene blocks used in this study are listed in Table S1. Full-length nsp1 was synthesized as a gene fragment from Integrated DNA technologies (IDT) and cloned into the EcoR1 restriction site of pGEX-6P-2 (GE healthcare) using in-fusion cloning (TakaraBio). Single primer-based mutagenesis was used to generate R124A/K125A, K164A/H165A, all N-terminal domain mutants (E36A/E37A, E55A/E56A/K57A, R99A, R119A/K120A) and the N-terminal cysteine/lysine containing nsp1 (nsp1 C-K) used in the fluorescence polarization assays was generated as mentioned above^48^. A T7 promoter upstream of the human β-globin 5’ UTR or the SARS-CoV-2 leader fused to a nano luciferase (HBB-nLuc and CoV-2 leader-nLuc, respectively) was synthesized by Twist Bioscience then subcloned by in-fusion cloning into an EcoRI site in a pUC57 destination vector. The leader sequence was derived from the SARS-CoV-2 isolate Wuhan-Hu-1 reference genome^49^. pCDNA4-3X-FLAG-Halo was used as a destination vector for mammalian cell transfection experiments for immunoprecipitations and GFP assays. CoV-2 nsp1 was subcloned into the NotI site of pCDNA4-3X-FLAG-Halo using in-fusion cloning. Single primer mutagenesis was used to generate R124A/K125A, K164A/H165A and all N-terminal mutants (E36A/E37A, E55A/E56A/K57A, R99A, R119A/K120A). Truncation mutants Δ118-180 and Δ1-117 were PCR amplified from the parental vector pCDNA4-3x-FLAG-Halo-nsp1 and cloned as described above. For the luciferase and RIP assays, HBB-nLuc was PCR amplified from the pUC57 vector (ThermoFisher) and cloned into AfeI and EcoRI digested pLJM1 vector (Addgene #19319) with InFusion. The CoV-2-leader nLuc sequence was synthesized by IDT and similarly cloned into the pLJM1 vector. All constructs were sequence verified by sanger sequencing.

### Protein expression and purification

WT or mutant nsp1 was expressed using BL21-CodonPlus (DE3)-RIPL cells (Agilent) using Overnight ExpressTM instant TB medium (Novagen). Cells were grown at 37°C to an OD600 of 0.6 then transferred to 18°C for 24 hr before being spun down at 4704 x g for 10 min. Cells were washed with PBS containing cOmpleteTM, EDTA-free protease inhibitor cocktail (Roche) and spun down once more. Wash solution was then decanted and pellets were resuspended in a buffer containing 500mM NaCl (Millipore sigma), 5mM MgCl_2_ (Millipore Sigma), 20mM HEPES (Millipore Sigma), 0.5% Triton x-100 (Sigma-Aldrich), 5% glycerol (Sigma-Aldrich) and 1mM Tris(2-Carboxyethyl)phosphine hydrochloride (Sigma-Aldrich) (Buffer A). The buffer pH was brought to 7.5 using concentrated hydrochloric acid solution (Sigma-Aldrich). The cell suspension was lysed at 4°C using a macrotip sonicator set at 80 amps with a 3 sec pulse and 17 sec rest for 12 min. Lysates were cleared by centrifuging at 50,000xg for 30 min at 4°C. The cleared lysate was incubated for 2 hr at 4°C on a rotating wheel with GST beads (Cytiva) that were pre-washed twice with buffer A. After incubation, beads were pelleted at 40 x g for 5 min at 4°C then washed 3x with Buffer A before resuspension in a buffer containing 250mM NaCl, 5mM MgCl_2_, 5% glycerol, 20mM HEPES pH 7.5, and 1mM TECEP (Buffer B). Beads were then loaded onto a polypropylene disposable column (Qiagen), and washed with 20 column volumes of Buffer B. After the final wash, beads were resuspended in one column volume of Buffer B. PreScission protease (Cytiva) was then added to the bead slurry and incubated on a rotating wheel overnight at 4°C. The elution was then collected and the beads were washed with an additional column volume of Buffer B. Both elutions were pooled and concentrated using a 10K Centriprep (EMD Millipore) before being loaded onto a HiLoad Superdex 200pg size exclusion column (Cytiva). Proteins eluted at expected masses as determined by protein standards and purity was determined using SDS-PAGE gel electrophoresis. Nsp1-containing fractions were then pooled and concentrated to 5 mg/ml and aliquots were stored at 80°C.

### *In vitro* transcription

RNA constructs were made by *in vitro* transcription with T7 RNA polymerase (New England Biolabs) as previously described^50^, with the following modifications. DNA templates were amplified from a plasmid containing the corresponding 5’ HBB UTR or CoV-2 leader sequence and the NanoLuc Luciferase coding sequence. Primers used for this amplification added a 60T sequence at the 3’ end to form a poly(A) tail after transcription. Transcription was then performed using gel-extracted PCR products and the T7 RNA polymerase New England Biolabs protocol. RNA was then capped using Vaccinia D1/D2 (Capping enzyme) (New England Biolabs) and 2’O-methylated using Vaccinia VP39 (2’O Methyltransferase) (New England Biolabs), then purified by phenol-chloroform extraction and ethanol precipitation.

### *In vitro* translation assays in rabbit reticulocyte lysate

*In vitro* translation reactions were performed using nuclease-treated rabbit reticulocyte lysate (RRL) (Promega) following the manufacturer’s protocol for non-radioactive luciferase reactions, with the following modifications. *In vitro* translation reactions were pre-incubated with the corresponding recombinant nsp1 variant for 10 min at 4°C before addition of 40nM of the corresponding mRNA. Reactions were then incubated at 30°C for 30 min, whereupon luciferase assays were performed using the NanoLuc luciferase assay kit (Promega). Luminescence was measured using the Spark multimode microplate reader (TECAN). Technical triplicate measurements were taken for each biological replicate and these technical triplicates were averaged to plot for a given biological replicate and normalized to GST.

### Preparation of HEK293T translation extracts

*In vitro* translation extracts were made from HEK293T cells using a previously described protocol^51^. Cells were scraped and collected by centrifugation for 2 minutes at 376 x g at 4°C. Cells were washed once with cold PBS (137mM NaCl, 2.7mM KCl, 100mM Na_2_HPO_4_, 2mM KH_2_PO_4_) then homogenized with an equal volume of freshly made cold hypotonic lysis buffer (10mM HEPES-KOH pH 7.6, 10mM KOAc, 0.5mM Mg(OAc)_2_, 5mM dithiothreitol (DTT), and 1 Complete EDTA-free Proteinase Inhibitor Cocktail tablet (Roche) per 10 ml of buffer). After hypotonic-induced swelling for 45 min on ice, cells were homogenized using a syringe attached to a 27G needle until 95% of cells burst as determined by trypan blue staining. Lysate was then centrifuged at 14,000 x g for 1 min at 4°C. The resulting supernatant was moved to a new tube, avoiding the top lipid layer. Lysate aliquots were quickly frozen with liquid nitrogen and stored at 80°C.

### *In vitro* translation assays

*In vitro* translation reactions were performed using HEK293T translation-competent cell lysate, as previously described, with modifications^50^. Translation reactions contained 50% HEK293T translation-competent cell extract, 2mM ATP, 0.42mM GTP, 7mM tris(2-carboxyethyl)phosphine, 28mM HEPES pH 7.5, 2mM creatine phosphate (Roche), 0.01 μg/μl creatine kinase (Roche), 2mM Mg(OAc)_2_, 60mM KOAc, 10μM amino acids (Promega), 0.21mM spermidine, 0.6mM putrescine and 0.8U/ml murine RNase inhibitor (NEB). Translation reactions were pre-incubated with the corresponding recombinant nsp1 variant for 10 min at 4°C before addition of the mRNA. 40nM of the corresponding RNA was then added to the reaction. Translation reactions were then incubated at 30°C for 30 min. Luciferase assays were then performed using the NanoLuc luciferase assay kit (Promega), following the manufacturer’s protocol. Luminescence was measured using the Spark multimode microplate reader (TECAN). Technical triplicate measurements were taken for each biological replicate and these technical triplicates were averaged to plot for a given biological replicate and normalized to GST.

### Primer extension assays

The 40μL reactions contained 50% HEK293T translation extract, 2mM ATP, 0.05mM GTP, 7mM tris(2-carboxyethyl)phosphine, 28mM HEPES pH 7.5, 2mM creatine phosphate (Roche), 0.01μg/μl creatine kinase (Roche), 2mM Mg(OAc)_2_ and 10μM of the amino acid mixture (Promega). Reactions were incubated with 5μM of the indicated nsp1 protein for 30 min on ice. 5μM of 5’-Guanylyl imidodiphosphate (GDPPNP) (Sigma-Aldrich) and 500μM of cycloheximide (CHX) (Sigma-Aldrich) was then added and incubated at RT for 10 min, whereupon 1μg of the HBB-nLuc mRNA was added and reactions were incubated at 30°C for 15 min. Reactions were stopped with the addition of 400μL of RNase free water and equal amounts of TRIzol reagent (ThermoFisher). RNA was extracted and resuspended in RNase free water then added to primer extension buffer containing 7mM MgOAc (Sigma-Aldrich), 100mM KOAc (Sigma-Aldrich), 50mM Tris-HCl, 1mM DTT, 1 μg/μL of SuperScript IV RT (Ambion), 550μM of DNTP mixture (ThermoFisher) and 500nM of Cy5 labeled primer (IDT). The reaction was incubated at 30°C for 15 minutes then ethanol precipitated. DNA pellets were washed twice with 70% ethanol, dried, and resuspended in 95% formamide solution containing 10mM EDTA. Samples were loaded onto a pre-run 6% UREA-PAGE gel then imaged using an Amersham Typhoon 5 laser-scanner (Cytiva).

### Cell culture and transfections

HEK293T cells were grown in Dulbecco’s modified Eagle’s medium (Gibco) supplemented with 10% fetal bovine serum (Peak Serum) at 37°C and 5% CO_2_. Transfections were carried out using PolyJet™ (SignaGen labs) according to the manufacturer’s DNA transfection protocol and cells were harvested 24 hr post transfection. For the GFP cotransfection experiments, 1×10^6^ cells were seeded into each well of a 6-well plate and transfected the next day with 100 ng of pCDNA-GFP and 900 ng of the indicated pCDNA4-3X-FLAG-Halo-nsp1 construct. For the luciferase assays, 4.5×10^4^ cells were seeded into each well of a 96-well plate and transfected the next day with 10 ng of HBB-nLuc or CoV2L-nLuc and 25 ng of the indicated C-terminally 3xFLAG tagged nsp1 (pCDNA4-nsp1-3xFLAG) construct. For the immunoprecipitation experiments, a 150 mm plate of cells were transfected with 25μg of the indicated pCDNA4-3X-FLAG-Halo-nsp1 construct. Cells were then harvested and washed with PBS (gibco) and snap frozen in liquid nitrogen and stored at −80°C until further processing.

### Luciferase assays

The media was aspirated off in the 96-well plate, and PBS was added onto the cells. The nanoluciferase luminescence was activated following the Nano-Glo Luciferase Assay System (Promega) protocol. The raw luminescence values were measured by TECAN SPARKCONTROL.

### RNA extraction and RT-qPCR

Cells were lysed in TRIzol reagent and RNA was extracted following the manufacturer’s protocol. The RNA was subsequently treated with TURBO DNase (ThermoFisher) and subjected to reverse transcription using AMV Reverse Transcripase (Promega) with a random 9-mer primer. cDNA was then used in qPCR following the iTaq Universal SYBR Green Supermix protocol (Bio-Rad laboratories) and gene-specific qPCR primers.

### Western blot analysis and nsp1 immunoprecipitation

For western blots on direct cell lysates, cell pellets were lysed in 2X laemmli buffer (BioRad laboratories) then homogenized using a microtip sonicator set at 20 amps, boiled for 10 min and resolved by SDS-PAGE. For immunoprecipitations, cell pellets were lysed in 1 mL of lysis buffer (Buffer A) containing 20mM Tris-KOH pH 7.5, 150mM KOAc, 5mM MgCl_2_, 5% glycerol, 1X HaltTM protease inhibitor cocktail (ThermoFisher), 1mM DTT (Danville Scientific) and 0.5% NP-40 (Danville Scientific) and placed on a rotating wheel for 30 min. Lysates were then passed through a 25G syringe needle ten times then spun at 21130 x g for 25 min to clear cell debris. Cleared lysates were combined with 20μL of ANTI-FLAG M2 (Sigma-Aldrich) magnetic beads that had been pre-washed 2x with Buffer A then incubated for 2 hr on a rotating wheel at 4°C. Beads were then washed 4x with Buffer A containing only 0.1% NP-40 and bound protein was eluted by 3 sequential incubations in PBS containing 0.1% triton x-100 (Sigma-Aldrich) and 0.5mg/ml of 3x FLAG peptide (Sigma-Aldrich) at 30°C for 20 min under vigorous shaking. The eluted protein was then precipitated using 100% trichloroacetic acid (Sigma-Aldrich) added drop wise to 20% of the final elution followed by vortexing and incubation at 4°C overnight prior to pelleting at 21130 x g for 20 min at 4°C. The protein pellet was washed with 100% cold acetone then dried at RT for 10 min, resuspended in 2X Laemmli buffer and boiled for 5 min prior to resolution by SDS-PAGE.

The following antibodies were used for western blotting: mouse anti-GFP (1:5000; Clontech 632381), rabbit anti-Vinculin (1:1000, Abcam GR268234-50), nd mouse antiFLAG M2 (1:1000, Sigma-Aldrich SLBT7654), rabbit anti-RPS2 (1:2000, Bethyl labs A303-794A-M), rabbit anti-RPS3 (1:500, Proteintech 11990-1-AP), rabbit anti-RPS24 (1:1000, Bethyl labs A303-842A-T), rabbit anti-RACK1 (1:1000, Bethyl labs A302-545A), HRP goat anti-mouse IgG (1:10,000 SouthernBiotech 1031-05) and HRP goat anti-rabbit IgG (1:10,000 SouthernBiotech 4030-05).

### RNA immunoprecipitation (RIP)

Cells were lysed in IP buffer (100 mM KCl, 0.1 mM EDTA, 20 mM HEPES-KOH pH 7.6, 0.4% NP-40, 10% glycerol; supplemented with fresh 1mM DTT, SUPERaseIN RNase Inhibitors (Ambion), and EDTA-free protease inhibitors (Roche)) then rotated at 4°C for 30 min and subsequently clarified by centrifugation at 21130 x g for 10 min at 4°C. The supernatant was brought up to 1 mL volume using IP buffer, and 90% of the sample was used for the RIP while 10% was saved as input samples. For each RIP, 20 μL of anti-FLAG M2 Magnetic beads (Sigma-Aldrich) were used, and the samples were rotated overnight at 4°C. The next day, samples were washed three times using 1 mL IP buffer with 5 min of rotation for each wash. The beads were then split into two fractions and one fraction was treated with TRIzolTM (ThermoFisher) for RNA extraction and the other was resuspended in Laemmli buffer (Bio-Rad laboratories) for western blot analysis. For the RNA extraction and RT-qPCR, the RIP samples were normalized to their respective input samples.

### TAMRA labeling of nsp1

Nsp1 C-K (3 mg/mL) was desalted using a 7K MWCO, 0.5 mL ZebaTM spin column (ThermoFisher) into a buffer containing 20mM HEPES-KOH pH 7.5, 120mM KOAc, 5mM Mg(OAc)_2_. A 1:5 mole ratio of protein to TAMRA maleimide, 6-isomer (lumiprobe Life science solutions) was used for each reaction. The reaction was gently mixed and placed at RT for 1 hr protected from light. The protein solution was then desalted once more to remove any free probe into a buffer containing 20mM HEPES-KOH pH 7.5, 120mM KOAc, 5mM Mg(OAc)_2_, 5% glycerol and 1mM TCEP. The dye:protein ratio for each labeled protein was between 0.84-0.91. Final protein concentration was determined by A280 and generally yielded a ~90 percent recovery.

### Ribosome purification

HeLa cell extract was prepared as described previously^52^. A frozen HeLa cell pellet was thawed and suspended with an equal volume of lysis buffer (20 mM HEPES pH 7.5, 10 mM KOAc, 1.8 mM Mg(OAc)_2_ and 1 mM DTT). After incubation on ice for 20 min, the cells were lysed with a Dounce homogenizer 150 times, followed by centrifugation twice at 1,200xg for 5 min. Supernatant was then aliquoted, flash-frozen with liquid nitrogen and stored at 80°C.

Crude 80S solutions were prepared from HeLa cell lysate using a previously described protocol^53^, with the following modifications. Lysate was loaded on 50% sucrose cushion prepared in Buffer A (20mM Tris pH 7.5, 2mM Mg(OAc)_2_, 150mM KCl), with the addition of 1mM dithiothreitol (DTT). Sucrose cushions were then centrifuged at 100,000rpm using an MLA-130 rotor (Beckman Coulter) for 60 min at 4°C. The resulting pellet was resuspended in cold Buffer A to homogeneity. Resuspended ribosome pellet was then centrifuged at 21130 x g for 10 min at 4°C to remove any remaining cell debris.

Supernatant was then aliquoted, flash-frozen with liquid nitrogen and stored at 80°C. 80S concentration was calculated as described previously^53,54^.

### Fluorescence polarization experiments

Binding experiments with TAMRA-labeled nsp1 were conducted using a Spark multimode microplate reader (TECAN). The final concentration of labeled nsp1 was limiting (5-10 nM), and the concentration of ribosomes was varied in each reaction. Binding reactions of 20μL were set up containing 20mM HEPES-KOH, 100mM KOAc, 5mM Mg(OAc)_2_, 5% glycerol, 0.2 mg/mL BSA (Ambion). Reactions were allowed to reach equilibrium at room temperature before the anisotropy was measured. Total fluorescence was also measured in order to account for quantum yield effects. The average change in anisotropy between free and bound nsp1 (and mean deviation) over three replicate experiments were: 0.078 ± 0.009. Multiple repeated measurements showed that equilibrium had been reached. Plotting and fitting the data to obtain Kd values was conducted assuming a “tight-binding” regime. This was based on recent publications where binding affinities of nsp1 to 40S subunits were determined^26^. Furthermore, saturation of binding was reached by 5 μM of ribosome, so the maximum anisotropy of the complex could be directly measured. The fraction bound was then calculated (incorporating quantum yield changes between the free and bound fluorophore) and fit to the solution of a quadratic equation describing an equilibrium reaction as previously reported^26,55,56^. The Kd values reported are the averages and the errors reported are the mean deviations.

## Acknowledgements

We thank David Morgens and Lucas Ferguson for helpful discussions throughout this project as well as for providing critical feedback on the manuscript. This project was funded by a COVID19 Excellence in Research Award from the Laboratory of Genomics Research to BAG, JHC and NTI and an Emergent Ventures COVID19 Fast Grant #2155 to BAG, who is also an investigator of the Howard Hughes Medical Institute. ML is funded by NSERC Predoctoral Fellowship PGSD3-516787-2018.

## References

1 Glaunsinger, B. A. Modulation of the Translational Landscape During Herpesvirus Infection. Annu Rev Virol 2, 311–333, doi:10.1146/annurev-virology-100114-054839 (2015).

2 Rivas, H. G., Schmaling, S. K. & Gaglia, M. M. Shutoff of Host Gene Expression in Influenza A Virus and Herpesviruses: Similar Mechanisms and Common Themes. Viruses 8, 102, doi:10.3390/v8040102 (2016).

3 Walsh, D. Manipulation of the host translation initiation complex eIF4F by DNA viruses. Biochem Soc Trans 38, 1511–1516, doi:10.1042/BST0381511 (2010).

4 Lloyd, R. E. Translational control by viral proteinases. Virus Res 119, 76–88, doi:10.1016/j.virusres.2005.10.016 (2006).

5 Abernathy, E. & Glaunsinger, B. Emerging roles for RNA degradation in viral replication and antiviral defense. Virology 479-480, 600–608, doi:10.1016/j.virol.2015.02.007 (2015).

6 Narayanan, K., Ramirez, S. I., Lokugamage, K. G. & Makino, S. Coronavirus nonstructural protein 1: Common and distinct functions in the regulation of host and viral gene expression. Virus Res 202, 89–100, doi:10.1016/j.virusres.2014.11.019 (2015).

7 Hartenian, E. et al. The molecular virology of coronaviruses. J Biol Chem 295, 12910–12934, doi:10.1074/jbc.REV120.013930 (2020).

8 Nakagawa, K. & Makino, S. Mechanisms of Coronavirus Nsp1-Mediated Control of Host and Viral Gene Expression. Cells 10, doi:10.3390/cells10020300 (2021).

9 Banerjee, A. K. et al. SARS-CoV-2 Disrupts Splicing, Translation, and Protein Trafficking to Suppress Host Defenses. Cell 183, 1325–1339 e1321, doi:10.1016/j.cell.2020.10.004 (2020).

10 Zhang, K. et al. Nsp1 protein of SARS-CoV-2 disrupts the mRNA export machinery to inhibit host gene expression. Sci Adv 7, doi:10.1126/sciadv.abe7386 (2021).

11 Hillen, H. S. et al. Structure of replicating SARS-CoV-2 polymerase. Nature 584, 154–156, doi:10.1038/s41586-020-2368-8 (2020).

12 Littler, D. R., Gully, B. S., Colson, R. N. & Rossjohn, J. Crystal Structure of the SARS-CoV-2 Non-structural Protein 9, Nsp9. iScience 23, 101258, doi:10.1016/j.isci.2020.101258 (2020).

13 Brockway, S. M. & Denison, M. R. Mutagenesis of the murine hepatitis virus nsp1-coding region identifies residues important for protein processing, viral RNA synthesis, and viral replication. Virology 340, 209–223, doi:10.1016/j.virol.2005.06.035 (2005).

14 Zust, R. et al. Coronavirus non-structural protein 1 is a major pathogenicity factor: implications for the rational design of coronavirus vaccines. PLoS Pathog 3, e109, doi:10.1371/journal.ppat.0030109 (2007).

15 Wathelet, M. G., Orr, M., Frieman, M. B. & Baric, R. S. Severe acute respiratory syndrome coronavirus evades antiviral signaling: role of nsp1 and rational design of an attenuated strain. J Virol 81, 11620–11633, doi:10.1128/JVI.00702-07 (2007).

16 Lei, L. et al. Attenuation of mouse hepatitis virus by deletion of the LLRKxGxKG region of Nsp1. PLoS One 8, e61166, doi:10.1371/journal.pone.0061166 (2013).

17 Kamitani, W. et al. Severe acute respiratory syndrome coronavirus nsp1 protein suppresses host gene expression by promoting host mRNA degradation. Proc Natl Acad Sci US A 103, 12885–12890, doi:10.1073/pnas.0603144103 (2006).

18 Narayanan, K. et al. Severe acute respiratory syndrome coronavirus nsp1 suppresses host gene expression, including that of type I interferon, in infected cells. J Virol 82, 4471–4479, doi:10.1128/JVI.02472-07 (2008).

19 Kamitani, W., Huang, C., Narayanan, K., Lokugamage, K. G. & Makino, S. A two-pronged strategy to suppress host protein synthesis by SARS coronavirus Nsp1 protein. Nat Struct Mol Biol 16, 1134–1140, doi:10.1038/nsmb.1680 (2009).

20 Huang, C. et al. SARS coronavirus nsp1 protein induces template-dependent endonucleolytic cleavage of mRNAs: viral mRNAs are resistant to nsp1-induced RNA cleavage. PLoSPathog 7, e1002433, doi:10.1371/journal.ppat.1002433 (2011).

21 Lokugamage, K. G., Narayanan, K., Huang, C. & Makino, S. Severe acute respiratory syndrome coronavirus protein nsp1 is a novel eukaryotic translation inhibitor that represses multiple steps of translation initiation. J Virol 86, 13598–13608, doi:10.1128/JVI.01958-12 (2012).

22 Thoms, M. et al. Structural basis for translational shutdown and immune evasion by the Nsp1 protein of SARS-CoV-2. Science 369, 1249–1255, doi:10.1126/science.abc8665 (2020).

23 Schubert, K. et al. Author Correction: SARS-CoV-2 Nsp1 binds the ribosomal mRNA channel to inhibit translation. Nat Struct Mol Biol 27, 1094, doi:10.1038/s41594-020-00533-x (2020).

24 Shi, M. et al. SARS-CoV-2 Nsp1 suppresses host but not viral translation through a bipartite mechanism. bioRxiv, doi:10.1101/2020.09.18.302901 (2020).

25 Yuan, S. et al. Nonstructural Protein 1 of SARS-CoV-2 Is a Potent Pathogenicity Factor Redirecting Host Protein Synthesis Machinery toward Viral RNA. Mol Cell 80, 1055–1066 e1056, doi:10.1016/j.molcel.2020.10.034 (2020).

26 Lapointe, C. P. et al. Dynamic competition between SARS-CoV-2 NSP1 and mRNA on the human ribosome inhibits translation initiation. Proc Natl Acad Sci U S A 118, doi:10.1073/pnas.2017715118 (2021).

27 Lei, X. et al. Activation and evasion of type I interferon responses by SARS-CoV-2. Nat Commun 11, 3810, doi:10.1038/s41467-020-17665-9 (2020).

28 Clark, L. K., Green, T. J. & Petit, C. M. Structure of Nonstructural Protein 1 from SARS-CoV-2. J Virol 95, doi:10.1128/JVI.02019-20 (2021).

29 Semper, C., Watanabe, N. & Savchenko, A. Structural characterization of nonstructural protein 1 from SARS-CoV-2. iScience 24, 101903, doi:10.1016/j.isci.2020.101903 (2021).

30 Tanaka, T., Kamitani, W., DeDiego, M. L., Enjuanes, L. & Matsuura, Y. Severe acute respiratory syndrome coronavirus nsp1 facilitates efficient propagation in cells through a specific translational shutoff of host mRNA. J Virol 86, 11128–11137, doi:10.1128/JVI.01700-12 (2012).

31 Tidu, A. et al. The viral protein NSP1 acts as a ribosome gatekeeper for shutting down host translation and fostering SARS-CoV-2 translation. RNA,doi:10.1261/rna.078121.120 (2020).

32 Benedetti, F. et al. Emerging of a SARS-CoV-2 viral strain with a deletion in nsp1. J Transl Med 18, 329, doi:10.1186/s12967-020-02507-5 (2020).

33 Lin, J. W. et al. Genomic monitoring of SARS-CoV-2 uncovers an Nsp1 deletion variant that modulates type I interferon response. Cell Host Microbe 29, 489–502 e488, doi:10.1016/j.chom.2021.01.015 (2021).

34 Min, Y. Q. et al. SARS-CoV-2 nsp1: Bioinformatics, Potential Structural and Functional Features, and Implications for Drug/Vaccine Designs. Front Microbiol 11, 587317, doi:10.3389/fmicb.2020.587317 (2020).

35 Gordon, D. E. et al. A SARS-CoV-2 protein interaction map reveals targets for drug repurposing. Nature 583, 459–468, doi:10.1038/s41586-020-2286-9 (2020).

36 Kozak, M. Circumstances and mechanisms of inhibition of translation by secondary structure in eucaryotic mRNAs. Mol Cell Biol 9, 5134–5142, doi:10.1128/mcb.9.11.5134 (1989).

37 Gaglia, M. M., Covarrubias, S., Wong, W. & Glaunsinger, B. A. A common strategy for host RNA degradation by divergent viruses. J Virol 86, 9527–9530, doi:10.1128/JVI.01230-12 (2012).

38 Sun, Y. et al. An R195K Mutation in the PA-X Protein Increases the Virulence and Transmission of Influenza A Virus in Mammalian Hosts. J Virol 94, doi:10.1128/JVI.01817-19 (2020).

39 Zhang, R. et al. The nsp1, nsp13, and M proteins contribute to the hepatotropism of murine coronavirus JHM.WU. J Virol 89, 3598–3609, doi:10.1128/JVI.03535-14 (2015).

40 Richner, J. M. et al. Global mRNA degradation during lytic gammaherpesvirus infection contributes to establishment of viral latency. PLoS Pathog 7, e1002150, doi:10.1371/journal.ppat.1002150 (2011).

41 Strelow, L. I. & Leib, D. A. Role of the virion host shutoff (vhs) of herpes simplex virus type 1 in latency and pathogenesis. J Virol 69, 6779–6786, doi:10.1128/JVI.69.11.6779-6786.1995 (1995).

42 Gaucherand, L. et al. The Influenza A Virus Endoribonuclease PA-X Usurps Host mRNA Processing Machinery to Limit Host Gene Expression. Cell Rep 27, 776–792 e777, doi:10.1016/j.celrep.2019.03.063 (2019).

43 Levene, R. E., Shrestha, S. D. & Gaglia, M. M. The influenza A virus host shutoff factor PA-X is rapidly turned over in a strain-specific manner. J Virol,doi:10.1128/JVI.02312-20 (2021).

44 Sarnow, P., Cevallos, R. C. & Jan, E. Takeover of host ribosomes by divergent IRES elements. Biochem Soc Trans 33, 1479–1482, doi:10.1042/BST20051479 (2005).

45 Yang, Y. & Wang, Z. IRES-mediated cap-independent translation, a path leading to hidden proteome. J Mol Cell Biol 11, 911–919, doi:10.1093/jmcb/mjz091 (2019).

46 Finkel, Y. et al. The coding capacity of SARS-CoV-2. Nature 589, 125–130, doi:10.1038/s41586-020-2739-1 (2021).

47 Sakuraba, S., Qilin, X., Kasahara, K., Iwakiri, J. & Kono, H. Modeling the SARS-CoV-2 nsp1-5’-UTR complex via extended ensemble simulations. bioRxiv, 2021.2002.2024.432807, doi:10.1101/2021.02.24.432807 (2021).

48 Mendez, A. S., Vogt, C., Bohne, J. & Glaunsinger, B. A. Site specific target binding controls RNA cleavage efficiency by the Kaposi’s sarcoma-associated herpesvirus endonuclease SOX. Nucleic Acids Res 46, 11968–11979, doi:10.1093/nar/gky932 (2018).

49 Chan, J. F. et al. Genomic characterization of the 2019 novel human-pathogenic coronavirus isolated from a patient with atypical pneumonia after visiting Wuhan. Emerg Microbes Infect 9, 221–236, doi:10.1080/22221751.2020.1719902 (2020).

50 Lee, A. S., Kranzusch, P. J. & Cate, J. H. eIF3 targets cell-proliferation messenger RNAs for translational activation or repression. Nature 522, 111–114, doi:10.1038/nature14267 (2015).

51 Rakotondrafara, A. M. & Hentze, M. W. An efficient factor-depleted mammalian in vitro translation system. Nat Protoc 6, 563–571, doi:10.1038/nprot.2011.314 (2011).

52 Khatter, H. et al. Purification, characterization and crystallization of the human 80S ribosome. Nucleic Acids Res 42, e49, doi:10.1093/nar/gkt1404 (2014).

53 Jan, C. H., Williams, C. C. & Weissman, J. S. Principles of ER cotranslational translocation revealed by proximity-specific ribosome profiling. Science 346, 1257521, doi:10.1126/science.1257521 (2014).

54 Algire, M. A. et al. Development and characterization of a reconstituted yeast translation initiation system. RNA 8, 382–397, doi:10.1017/s1355838202029527 (2002).

55 Fraser, C. S., Berry, K. E., Hershey, J. W. & Doudna, J. A. eIF3j is located in the decoding center of the human 40S ribosomal subunit. Mol Cell 26, 811–819, doi:10.1016/j.molcel.2007.05.019 (2007).

56 Fraser, C. S., Hershey, J. W. & Doudna, J. A. The pathway of hepatitis C virus mRNA recruitment to the human ribosome. Nat Struct Mol Biol 16, 397–404, doi:10.1038/nsmb.1572 (2009).

