## Supplemental figures for "The N-terminal and central domains of CoV-2 nsp1 play key functional roles in suppression of cellular gene expression and preservation of viral gene expression"

Figure S1.

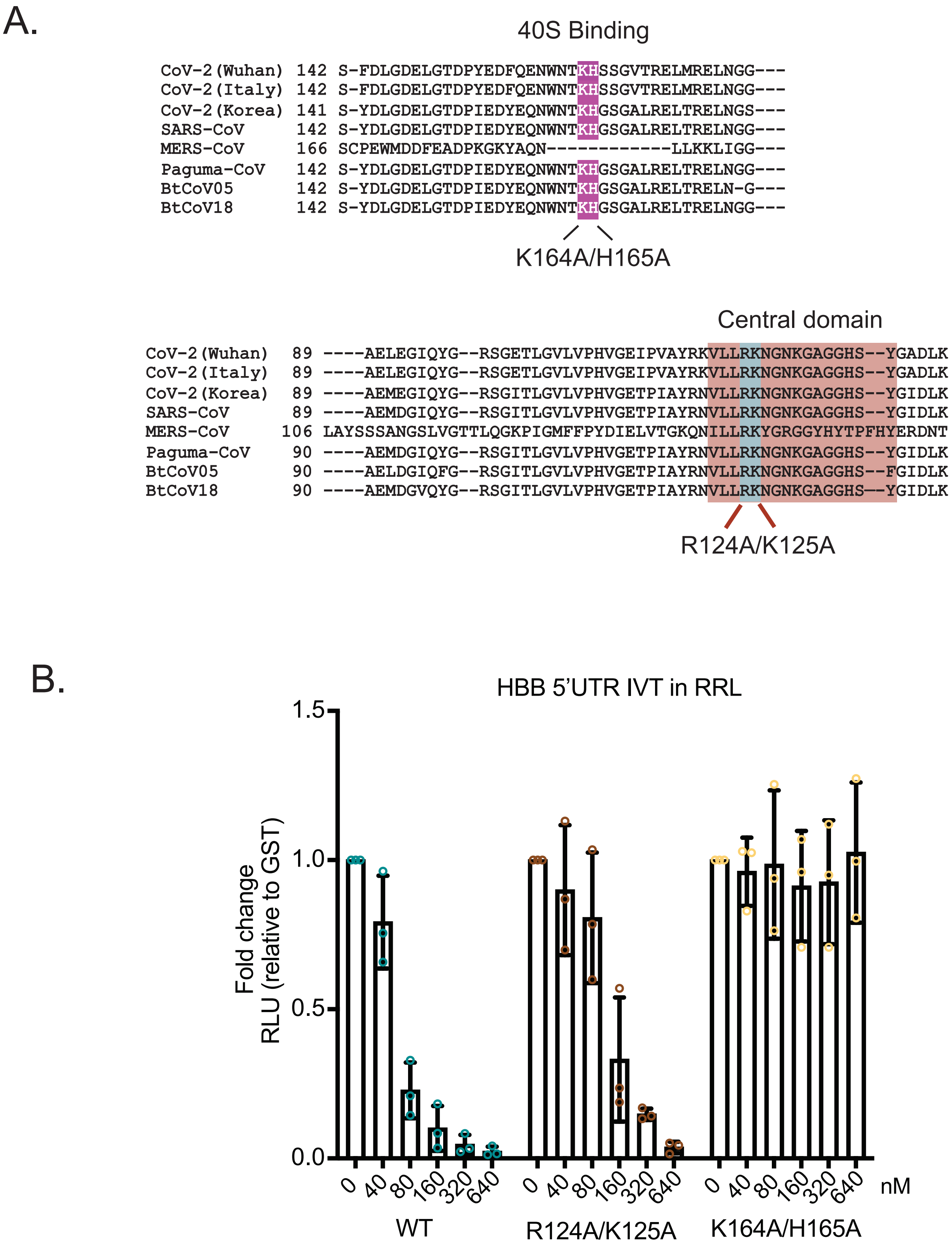


**Figure S1. Sequence alignment of the nsp1 C-terminal and central domains and RRL translation assay.** (A) The purple box highlights conservation of the K164 and H165 residues and the red box shows the central domain containing conserved residues R124/K125 (blue) involved in mRNA destabilization across 8 betacoronavirus nsp1 proteins. (B) HBB-nLuc reporter RNA was incubated with rabbit reticulocyte lysate translation extracts alone or in the presence of increasing concentrations of purified WT, R124A/K125A or K164A/H165A nsp1. Translation of the reporter was then evaluated by luciferase assay and normalized to a GST protein control. Data represent a total of 3 biological replicates.

Figure S2.

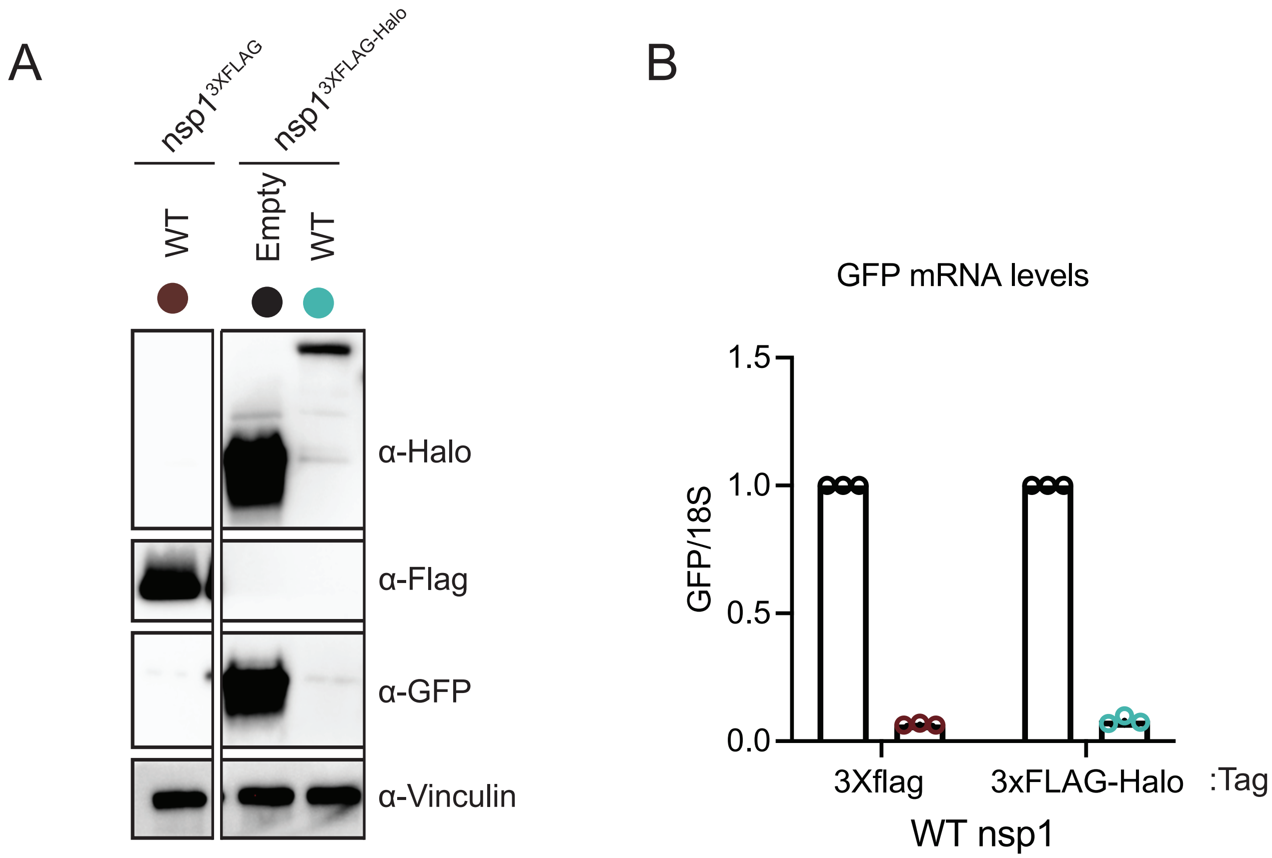


**Figure S2. Western blot and RT-qPCR analysis of 3xFLAG-Halo-tagged nsp1 and 3xFLAG tagged nsp1**. HEK293T cells were co-transfected with plasmids expressing GFP and either nsp1-3XFLAG or nsp1-3xFLAG-Halo and lysates were harvested for either protein or RNA. (A) Protein levels were measured by western blotting with antibodies against FLAG or Halo to detect nsp1 and antibodies against GFP as a marker of nsp1 host shutoff activity. Vinculin was used as a loading control. (B) The impact of each tagged version of nsp1 on GFP mRNA levels was determined by RT-qPCR and normalized to 18S rRNA, with the level of GFP mRNA in cells lacking nsp1 then set to 1. Each dot represents an independent experiment.

Figure S3.

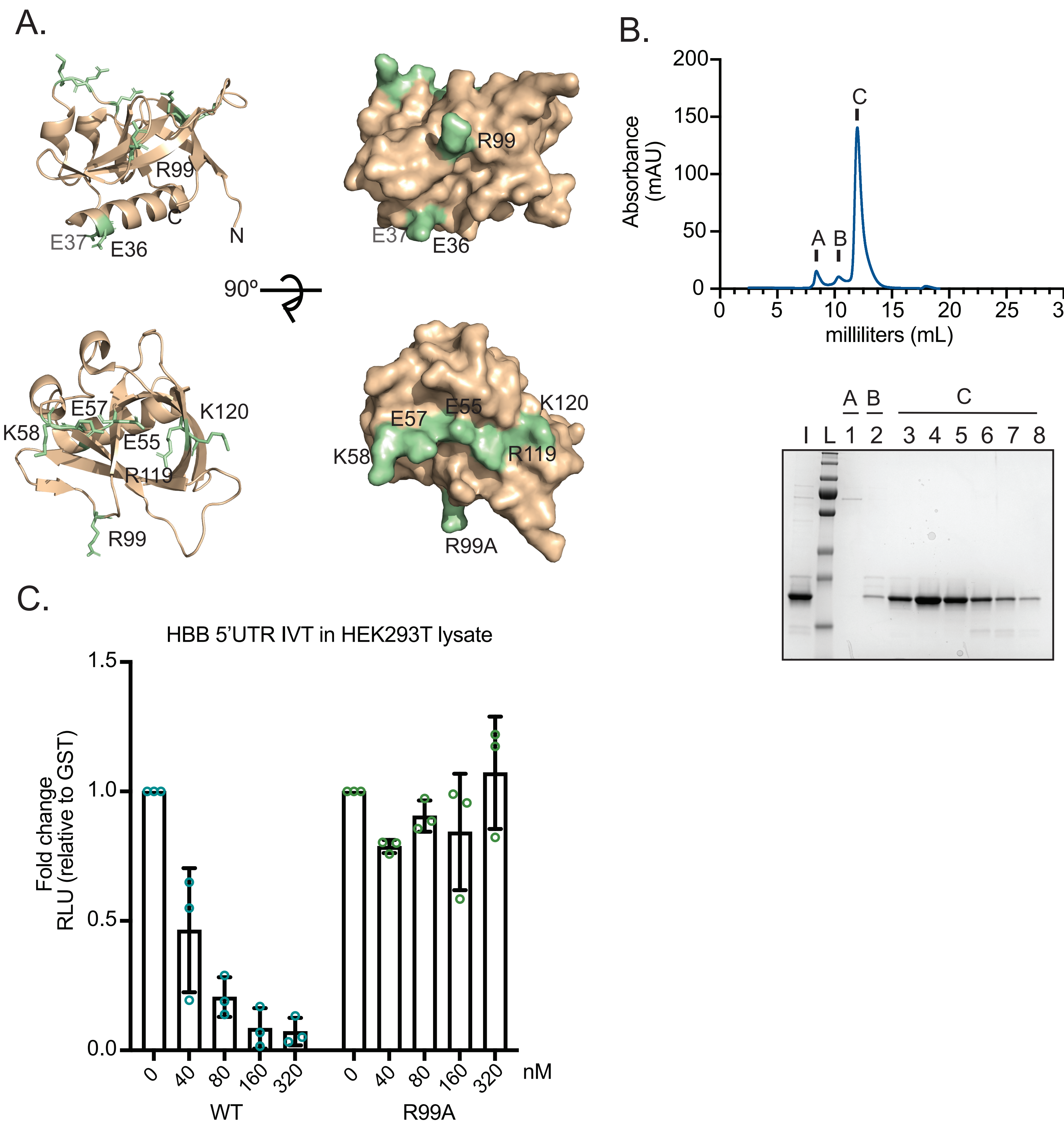


**Figure S3. Mutations to the N-terminal globular domain of nsp1**. (A) Structure of the nsp1 N-terminal domain (PDB: 7K7P) (Clark et al. 2021). Residues selected for mutation are highlighted in green. (B) Size exclusion run of the purified nsp1 R99A protein. (I) indicates input sample and peaks A and B represent higher molecular weight contaminants. Peak C corresponds to the expected molecular weight of the nsp1 R99A mutant. (L) corresponds to the prestained ladder. (C) HEK293T translation extracts were used to monitor the effect of increasing concentrations of WT nsp1 versus the R99A mutant on translation of HBB-nLuc reporter, as measured by luciferase assay. The data represent 3 independent experiments.

Figure S4.

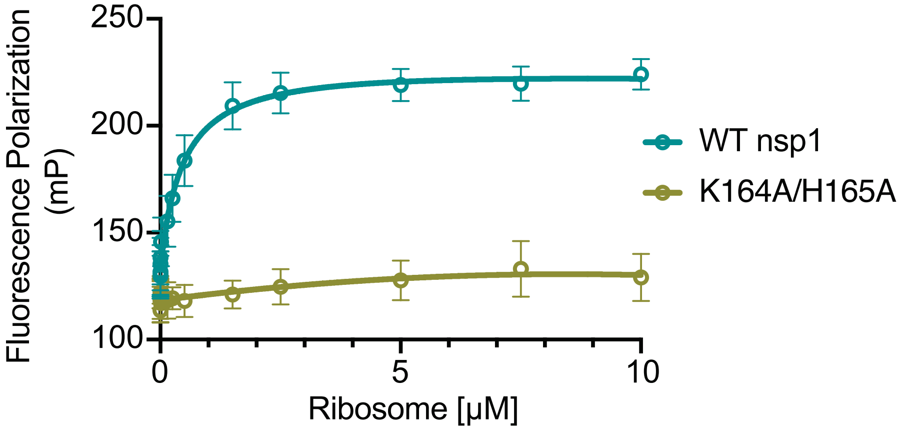


**Figure S4. Equilibrium binding of fluorescently labeled K164A/H165A nsp1**. Binding experiments were performed with nsp1 WT (blue circles) and K164A /H165A (green circles) and purified ribosomes. Raw millipolarization units are shown on the Y axis (mP).

Figure S5.

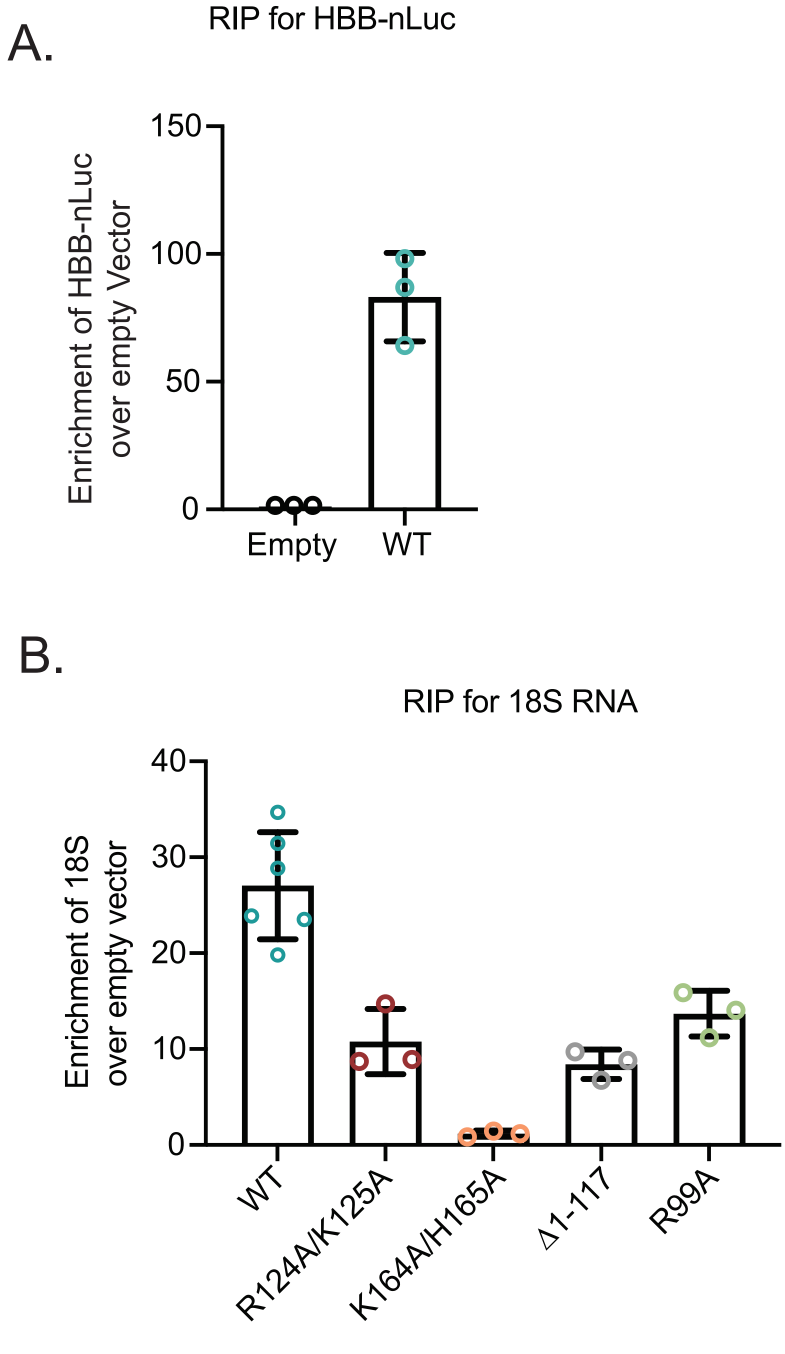


**Figure S5. Binding of HBB-nLuc mRNA and 18S rRNA to WT and/or mutant nsp1.** (A) RIP data showing the enrichment value of HBB-nLuc mRNA for WT nsp1 compared to empty vector, using the data from the experiments in Figure 4C. (B) RT-qPCR was performed to quantify 18S levels in the RIP experiment shown in Figure 4C, with the RNA values then normalized to the RNA values obtained from the empty vector control.

Figure S6.

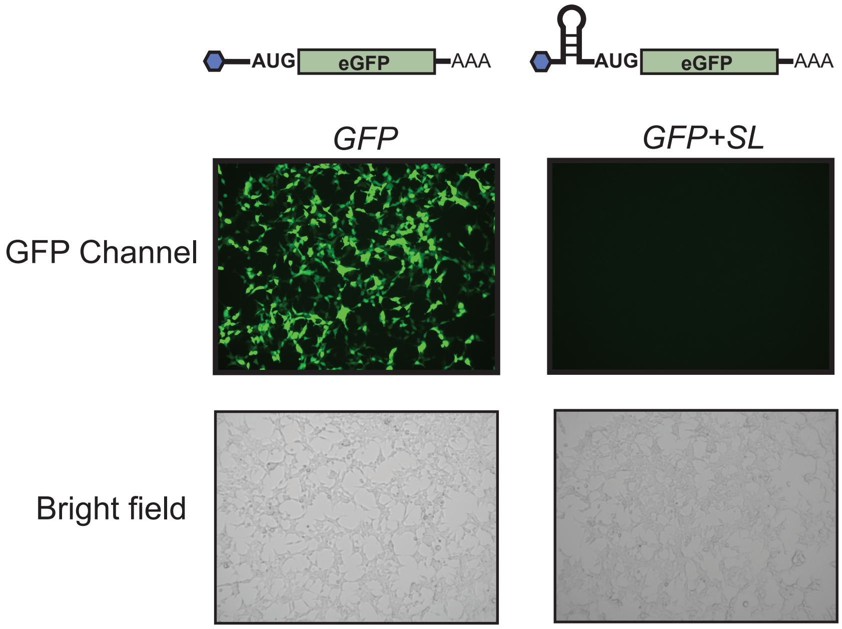


**Figure S6. Placement of a 40S blocking stem loop proximal to the 5' cap on the GFP reporter prevents its translation.** Plasmids encoding GFP mRNA or GFP mRNA containing a cap-proximal stem loop structure (*GFP+SL*) were transfected into HEK293T cells, and GFP fluorescence from each cell was assessed by fluorescence microscopy. Bright field images show cell density; all images were taken at 10X magnification.

Figure S7.

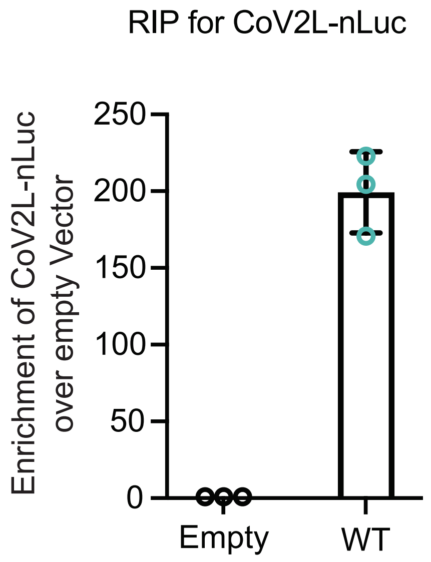


**Figure S7. Binding of CoV2L-nLuc mRNA to WT nsp1.** RIP data showing the enrichment value of CoV2L-nLuc mRNA for WT nsp1 compared to empty vector, using the data from the experiments in Figure 5B.

**Table S1. Oligos used in this study.**

| | | |
|---|---|---|
| **RT-qPCR primers** | | |
| Target gene | Forward primer | Reverse primer |
| GFP | GAACCGCATCGAGCTGAA | TGCTTGTCGGCCATGATATAG |
| Nanoluciferase | GGAGGTGTGTCCAGTTTGTT | ATGTCGATCTTCAGCCCATTT |
| 18S | GTAACCCGTTGAACCCCATT | CCATCCAATCGGTAGTAGCG |
| **Primer extension oligo** | | |
| Target gene | Sequence | |
| Nanoluciferase | CGCCAGAATGCGTTCGCACAGCCGCCAGCCGGTC | |
| **Oligos used for cloning** | | |
| Name | Sequence | Description |
| pGEX-CoV2nsp1_Fw | GGATCCCCAGGAATTGAGAGCCTTGTCCCTGGTTTC | Forward primer to amplify CoV-2 nsp1 and clone into pGEX vector via InFusion reaction for protein expression |
| pGEX-CoV2nsp1_Rv | AGTCACGATGCGGCCTTACCCTCCGTTAAGCTCACGC | Reverse primer to amplify CoV-2 nsp1 and clone into pGEX vector via InFusion reaction for protein expression |
| pCDNA4-3xFLAG-HaloTEV-CoV2_Fw | GACAAGGGGGCGGCCGCAGAAATCGGTACTGGCTTTCC | Forward primer to amplify CoV-2 nsp1 and generate N-terminally 3xFLAG-HaloTEV nsp1 via InFusion reaction |
| pCDNA4-3xFLAG-HaloTEV-CoV2_Rv | TAGACTCGAGCGGCCCTACCCTCCGTTAAGCTCACG | Reverse primer to amplify CoV-2 nsp1 and generate N-terminally 3xFLAG-HaloTEV nsp1 via InFusion reaction |
| pCDNA4-CoV2-FLAG_Fw | cacagtggcggccgcATGgagagccttgtccctggtttc | Forward primer to amplify CoV-2 nsp1 and generate C-terminally 3xFLAG tagged nsp1 via restriction enzyme cloning |

| | | |
|---|---|---|
| pCDNA4-CoV2-FLAG_Rv | ctcgagcggccgccaccctccgttaagctcacgcatgag | Reverse primer to amplify CoV-2 nsp1 and generate C-terminally 3xFLAG tagged nsp1 via restriction enzyme cloning |
| pCDNA4-CoV2-delta118-180_Fw | GACAAGGGGGCGGCCGCAGAAATCGGTACTGGCTTTCC | Forward primer to amplify the N-terminal half of CoV-2 nsp1 and generate CoV-2 nsp1 Δ 118-180 via InFusion reaction |
| pCDNA4-CoV2-delta118-180_Rv | TAGACTCGAGCGGCCCTAAGCCACTGGTATTTCGCCC | Reverse primer to amplify the N-terminal half of CoV-2 nsp1 and generate CoV-2 nsp1 Δ 118-180 via InFusion reaction |
| pCDNA4-CoV2-delta1-117_Fw | GACAAGGGGGCGGCCGCAGAAATCGGTACTGGCTTTCC | Forward primer to amplify the C-terminal half of CoV-2 nsp1 and generate CoV-2 nsp1 Δ 1-117 via InFusion reaction |
| pCDNA4-CoV2-delta1-117_Rv | TAGACTCGAGCGGCCCTACCCTCCGTTAAGCTCACG | Reverse primer to amplify the C-terminal half of CoV-2 nsp1 and generate CoV-2 nsp1 Δ 1-117 via InFusion reaction |
| CoV-2 nsp1 R124A/K125A | cgcaaggttcttcttGCtGCgaacggtaataaagga | Mutagenesis primers to generate R124A/K125A mutant |
| CoV-2 nsp1 K164A/H165A | gaaaactggaacactGCaGCtagcagtggtgttacc | Mutagenesis primers to generate K164A/H165A mutant |
| CoV-2 nsp1 E36A/E37A | GGCTTTGGAGACTCCGTGGCAGCAGTCTTATCAGAGGCAC | Mutagenesis primers to generate E36A/E37A mutant |
| CoV-2 nsp1 E55A/E57A/K58A | GGCACTTGTGGCTTAGTAGCAGTTGCAGCGGGCGTTTTGCCTCAAC | Mutagenesis primers to generate E55A/E57A/K58A mutant |
| CoV-2 nsp1 R99A | CGAAGGCATTCAGTACGGTGCAAGTGGTGAGACACTTGG | Mutagenesis primers to generate R99A mutant |
| CoV-2 nsp1 R119A/K120A | GAAATACCAGTGGCTTACGCAGCGGTTCTTCTTCGTAAGAAC | Mutagenesis primers to generate R119A/K120A mutant |
| CoV-2 nsp1 Δ 122-130 | CAGTGGCTTACCGCAAGGTTGCTGGTGGCCATAGTTACG | Mutagenesis primers to generate Δ 122-130 mutant |
| CoV-2 nsp1 G-linker | CAGTGGCTTACCGCAAGGTTGGAGGAGGAGGAAGCGGAGGAGGAGGAGCTGGTGGCCATAGTTAC | Mutagenesis primers to insert Glycine linker into the central domain |

| | | |
|---|---|---|
| | G | |
| N-terminal cysteine/lysine CoV-2 nsp1 | GAAGTTCTGTTCCAGGGGCCCT GTAAAGAGAGCCTTGTCCCTGG | Primers used to add cysteine and lysine to CoV-2 nsp1 for fluorescence polarization |
| pJP-HBB-nLuc_Fw | gtcagatccgctagcgctACATTTG CTTCTGAC | Forward primer to amplify HBB-nLuc and clone into pJP vector via InFusion reaction |
| pJP-HBB-nLuc_Rv | ttCTCTAGAGATATCttacgcca gaatgcgttcgca | Reverse primer to amplify HBB-nLuc and clone into pJP vector via InFusion reaction |
| CoV-2 leader geneblock | AAATGGACTATCATATGCC AAGTACGCCCCCTATTGAC GTCAATGACGGTAAATGGC CCGCCTGGCATTATGCCCA GTACATGACCTTATGGGAC TTTCCTACTTGGCAGTACA TCTACGTATTAGTCATCGC TATTACCATGGTGATGCGG TTTTGGCAGTACATCAATG GGCGTGGATAGCGGTTTG ACTCACGGGGATTTCCAAG TCTCCACCCCATTGACGTC AATGGGAGTTTGTTTTGGC ACCAAAATCAACGGGACTT TCCAAAATGTCGTAACAAC TCCGCCCCATTGACGCAAA TGGGCGGTAGGCGTGTAC GGTGGGAGGTCTATATAAG CAGAGCTGGTTTAGTGAAC CGACCTTCCCAGGTAACAA ACCAACCAACTTTCGATCT CTTGTAGATCTGTTCTCTAA ACGAACATGGTCTTCACAC TCGAAGATTTCGTTGGGGA CTGGCGACAGACAGCCGG CTACAACCTGGACCAAGTC CTTGAACAGGGAGGTGTGT CCAGTTTGTTTCAGAATCT CGGGGTGTCCGTAACTCC GATCCAAAGGATTGTCCTG AGCGGTGAAAATGGGCTG AAGATCGACATCCATGTCA TCATCCCGTATGAAGGTCT GAGCGGCGACCAAATGGG CCAGATCGAAAAAATTTTTA AGGTGGTGTACCCTGTGG | CoV-2 leader geneblock was synthesized by IDT and cloned into pJP vector via InFusion reaction |

| | | |
|---|---|---|
| | ATGATCATCACTTTAAGGT<br>GATCCTGCACTATGGCACA<br>CTGGTAATCGACGGGGTTA<br>CGCCGAACATGATCGACTA<br>TTTCGGACGGCCGTATGAA<br>GGCATCGCCGTGTTCGAC<br>GGCAAAAAGATCACTGTAA<br>CAGGGACCCTGTGGAACG<br>GCAACAAAATTATCGACGA<br>GCGCCTGATCAACCCCGA<br>CGGCTCCCTGCTGTTCCGA<br>GTAACCATCAACGGAGTGA<br>CCGGCTGGCGGCTGTGCG<br>AACGCATTCTGGCGTAGGA<br>ATTCTCGACCTCGA | |
| T7+HBB5'UTR+Koz+Nluc_F | taatacgactcactataggACATTTG<br>CTTCTGACACAACTGTGTTC<br>ACTAGCAACCTCAAACAGAC<br>ACCGCCACCATGGTCTTC | Forward primer to generate HBB-nLuc template for in-vitro transcription |
| NLuc_R_60T | TTTTTTTTTTTTTTTTTTTTTT<br>TTTTTTTTTTTTTTTTTTTTTT<br>TTTTTTTTTTTTTTTTTTACGC<br>CAGAATGCGTTCGCAC | Reverse primer to generate HBB-nLuc template for in-vitro transcription |
